# Dissecting peri-implantation development using cultured human embryos and embryo-like assembloids

**DOI:** 10.1101/2023.06.15.545180

**Authors:** Zongyong Ai, Baohua Niu, Yu Yin, Lifeng Xiang, Gaohui Shi, Kui Duan, Sile Wang, Yingjie Hu, Chi Zhang, Chengting Zhang, Lujuan Rong, Ruize Kong, Tingwei Chen, Yixin Guo, Wanlu Liu, Nan Li, Shumei Zhao, Xiaoqing Zhu, Xuancheng Mai, Yonggang Li, Ze Wu, Yi Zheng, Jianping Fu, Weizhi Ji, Tianqing Li

## Abstract

Studies of cultured embryos have provided insights into human peri-implantation development. However, detailed knowledge of peri-implantation lineage developments as well as underlying mechanisms remain obscure. Using 3D-cultured human embryos, herein we report a complete cell atlas of the early post-implantation lineages and decipher cellular composition and gene signatures of the epiblast and hypoblast derivatives. In addition, we develop an embryo-like assembloid (E-assembloid) by assembling naive hESCs and extraembryonic cells. Using human embryos and E-assembloids, we reveal that WNT, BMP and Nodal signaling synergistically, but functionally differently, orchestrate human peri-implantation lineage development. Specially, we dissect mechanisms underlying extraembryonic mesoderm and extraembryonic endoderm specifications. Finally, an improved E-assembloid is developed to recapitulate the epiblast and hypoblast developments and tissue architectures in the pre-gastrulation human embryo. Our findings provide insights of the human peri-implantation development, and the E-assembloid offers a useful model to disentangle cellular behaviors and signaling interactions that drive human embryogenesis.

## Introduction

In humans, the blastocyst, a hollow round structure comprising ∼200 cells, implants into the uterine wall to initiate post-implantation development at embryonic day 6-7 (E6-7). In the late blastocyst, the inner cell mass (ICM) segregates into the hypoblast (HB), which gives rise to the yolk sac, and the epiblast (EPI), which differentiates into the three definitive germ layers to form the embryo proper. Despite their critical roles in human development, we still have very limited knowledge of post-implantation development of the HB and EPI lineages. Recently, we and others have successfully developed *in vitro* human embryo culture systems, extending growth of human blastocysts towards the pre-gastrulation stage ^1–4^. However, the precise cell atlas, lineage specification and developmental signals during the peri-implantation human development, especially those related to the HB and EPI derivatives, remain obscure ^5^.

The use of human embryos for investigating peri-implantation development is fraught with ethical concerns and hampered by technical barriers, such as genetic manipulation in specific lineages and reproducible embryo development ^5^. Stem cell-based embryo models provide a more practical alternative in place of human embryos to decode early human development. Previous studies used primed human pluripotent stem cells (hPSCs) to model some embryonic phenotypes associated with post-implantation EPI development such as embryoid body, micropatterned colony, asymmetric human epiblast models (3D epiblast model and post-attached embryoid), gastruloid, sequential somite-like structures (Axioloid or Segmentoid) and amniotic sac embryoid ^6–16^. However, these embryoids could not recapitulate the specification and development of the extraembryonic mesoderm (ExM) and HB lineages, only mimicked some aspects of human postimplantation EPI development such as anteroposterior symmetry breaking, anterior-posterior axial patterning, somitogenesis, dorsal–ventral patterning and amniogenesis. Although several recent reports showed that human blastocyst-like structures (blastoids) are generated from naive hPSCs or human extended pluripotent stem cells (hEPSCs) or primed-to-naïve intermediates or by reprogramming fibroblasts, these blastoids showed a poor developmental potential through implantation ^17–23^. To our knowledge, some vital 3D tissue structures in human peri-implantation embryos, such as the amniotic cavity and yolk sac, are not well recapitulated in extended cultured human blastoids.

To fill the knowledge gap of human peri-implantation development, herein we provided a complete cell atlas of early post-implantation embryonic lineages in 3D cultured human blastocysts. Furthermore, we successfully created a human embryo-like assembloid (E-assembloid) that recapitulates developmental landmarks, 3D cellular organization and lineage composition of human peri-implantation embryo, thus establishing a novel embryo model for studying human early post-implantation development. Using human embryos and E-assembloids, our study unravels the transcriptional states of HB and EPI derivatives and functional roles of WNT, BMP and Nodal signaling in mediating their lineage specifications.

## Results

### Single-cell transcriptional profiling of 3D-cultured human embryos

Although some cell types in the peri-implantation human embryo have recently been characterized ^3, 4, 24^, a complete cell atlas of the post-implantation HB and EPI derivatives and their corresponding gene expression profiles, especially for amniotic epithelium (AME), extraembryonic mesoderm (ExM), visceral/yolk endoderm (VE/YE) and anterior visceral endoderm (AVE), are still lacking. To address this knowledge gap, we first generated a complete cell atlas for 3D-cultured human embryos ^4^ from embryonic day (E10-14) (Supplementary information, Fig. S1a) using the 10X Genomics platform. Different cell types, including trophoblast (TrB), EPI, ExM and extraembryonic endoderm (XEN, an HB derivative), emerged successively between E10-14 in the single-cell RNA-sequencing (scRNA-seq) data (Fig. 1a, b; Supplementary information, Fig. S1b). To focus on EPI and HB lineage developments, we removed TrB cells from the scRNA-seq data. Remaining cells were subclustered into 11 different populations, including post-implantation early and late EPI (PostE−EPI and PostL−EPI), undefined cells (UC), AME1 and 2, primitive streak 1 and 2 (PS1 and 2), ExM1 and 2, VE/YE and AVE (Fig. 1c, d; Supplementary information, Fig. S1c).

**Fig. 1.**
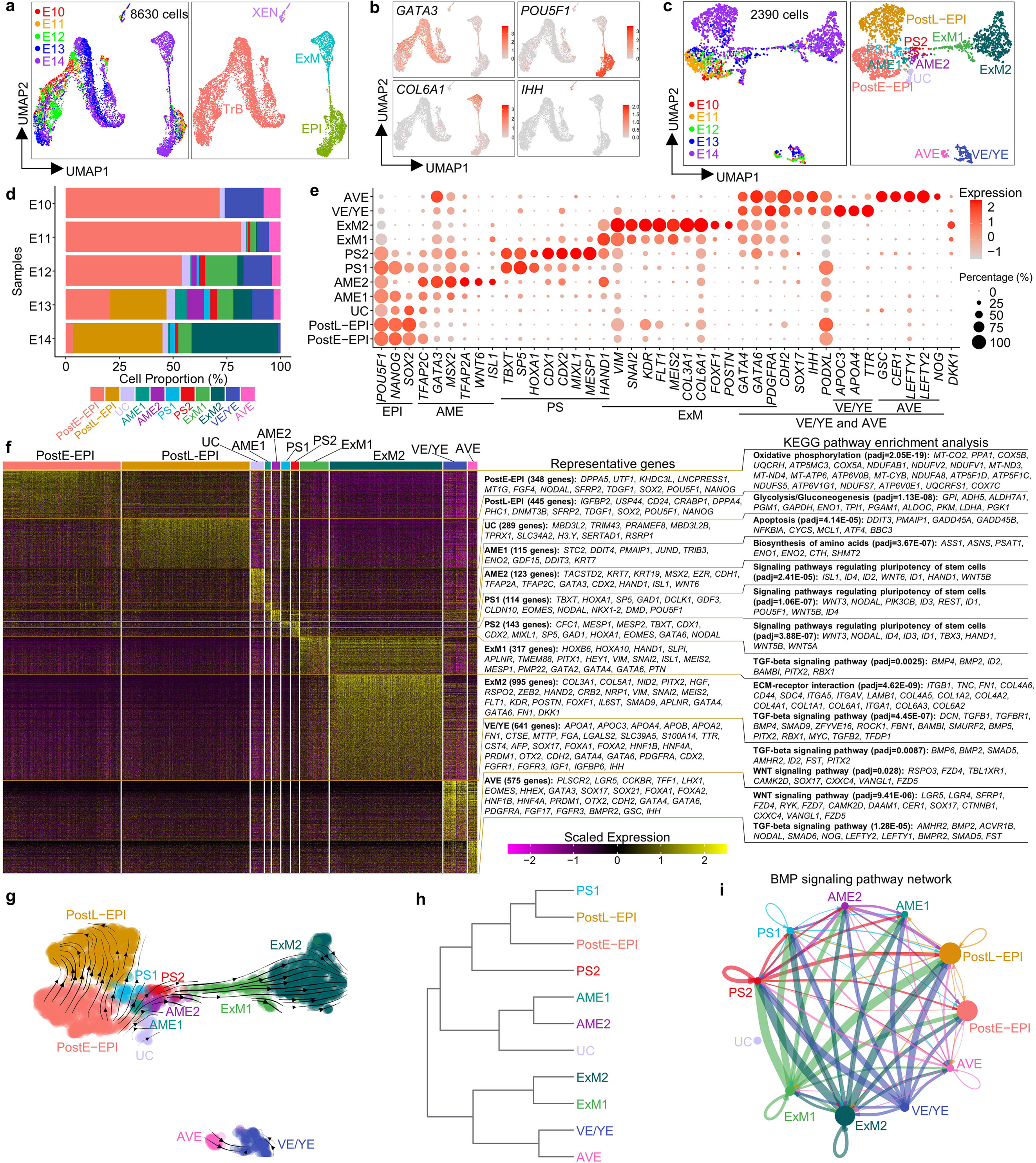
Single-cell transcriptome analysis of 3D-cultured human post-implantation embryos. The single-cell RNA sequencing (scRNA-seq) data are from 10x Genomics platform. **a** Uniform Manifold Approximation and Projection (UMAP) visualization of single-cell transcriptome datasets from 3D-cultured human post-implantation embryos at different embryonic days. **b** UMAP plots of genes that were differentially expressed in four different populations. **c** UMAP visualization of all cell types after excluding trophoblast subpopulation. **d** Proportions of indicated subtypes of cells in extended cultured human embryos at different developmental time points according to the results of scRNA-seq data. **e** Dot plots of candidate genes specific for cell subtypes. **f** Heatmap of differentially expressed genes (DEGs) among different cell subtypes displayed in **c**. Representative genes (left) and KEGG pathway enrichment analysis (right) are shown. P adj of Wilcoxon’s rank-sum test < 0.05, log_2_ (FC)□>□0.25 and expressed in >25% of cells of the given cluster. **g** RNA velocity vectors projected onto the UMAP-based embeddings of the single-cell transcriptome dataset from human embryos shown in **c**. **h** The hierarchical clustering tree of different cell subpopulations. **i** The inferred BMP signaling pathway network. Circle sizes are proportional to the number of cells in each subpopulation and line weight represents the communication probability. E, embryonic day; TrB, trophoblast; XEN, extraembryonic endoderm; ExM, extraembryonic mesoderm; EPI, epiblast; PostE−EPI and PostL−EPI, post-implantation early and late EPI; UC, undefined cell-type; AME, amniotic epithelium; PS, primitive streak; VE/YE, visceral/yolk sac endoderm; AVE, anterior visceral endoderm. See also Supplementary information, Fig. S1.

We next analyzed gene expression signatures of these lineages and identified molecular markers useful for discerning different cell types in the peri-implantation human development. As expected, both PostE−EPI and PostL−EPI express core pluripotency genes *POU5F1*, *NANOG* and *SOX2*. PostE−EPI expresses greater levels of naive genes *UTF1*, *FGF4* and *NODAL*, whereas PostL−EPI shows higher expression of primed genes *IGFBP2*, *USP44*, *CD24* and *DNMT3B* (Fig. 1e, f). KEGG (Kyoto Encyclopedia of Genes and Genomes) pathway analysis revealed that PostE−EPI and PostL−EPI are enriched with genes for ‘oxidative phosphorylation’ and ‘glycolysis’, respectively (Fig. 1f), suggesting a metabolic shift during EPI development. We also examined expression of genes associated with the formative pluripotency state ^25, 26^; there is greater expression of *DPPA2* and *GDF3*, but lower or similar expression of *SOX11*, *OTX2* and *ZNF* genes in PostE−EPI, compared to in PostL−EPI (Supplementary information, Fig. S1d), suggesting that there are differences between formative hPSCs and embryonic epiblast in gene expression profiles. The UC cluster weakly expresses pluripotency genes and is enriched with genes for ‘apoptosis’, suggesting their apoptotic EPI identity (Fig. 1e, f). Compared with the AME2, the AME1 shows lower expression of AME markers but higher expression of pluripotency genes; compared with the EPI, the AME1 shows lower expression of pluripotency markers but higher expression of amnion genes, indicative of a transitional AME state (Fig. 1e, f). RNA velocity map and clustering further reveal the similar identities of AME1 and AME2 (Fig. 1g, h). PS1 and PS2 share early primitive streak genes *TBXT*, *SP5*, *HOXA1*, *EOMES* and *NODAL*, with PS2 highly expressing *CDX1*, *CDX2*, *MIXL1*, *MESP1*, *HAND1* and *GATA6* (Fig. 1e, f). ExM1 and ExM2 upregulate epithelial to mesenchymal transition genes *VIM* and *SNAI2*; in addition to *KDR* ^4^ and *COL6A1* ^27^, we identified *FLT1*, *MEIS2*, *COL3A1*, *NID2* and *FOXF1* as potential markers of ExM (Fig. 1e; Supplementary information, Fig. S1e). Unexpectedly, *POSTN*, a specific AVE marker in marmoset ^28^, is exclusively expressed in the ExM2 (Fig. 1e; Supplementary information, Fig. S1e). Expressions of *GATA4*, *GATA6*, *PDGFRA*, *FN1* and *COL4A1* are all upregulated in the ExM (ExM1 and ExM2) and XEN (VE/YE and AVE), in contrast to *APOA1* ^29^, *IHH* ^30^, *OTX2* ^29^, *FOXA1* ^27^, *FOXA2*, *HNF4A* and *SOX17*, which are specific to XEN (Fig. 1e; Supplementary information, Fig. S1e). *AFP*, *TTR*, *APOA4*, *APOC3* and *MTTP* show upregulated expression in VE/YE. Conversely, *LHX1*, *HHEX*, *EOMES*, *GSC*, *CER1*, *LEFTY1*, *LEFTY2* and *NOG* are mainly expressed in AVE (Fig. 1e; Supplementary information, Fig. S1e), suggesting that human AVE is conserved to secret inhibitors of WNT, BMP and Nodal signaling.

We further performed immunostaining for E13 human embryos and confirmed that XEN cells are positive for GATA6, SOX17, FOXA1, PODXL1, N-Cadherin and OTX2, and ExM cells are positive for GATA6, KDR, SNAIL and COL3A1 but negative for SOX17, FOXA1, PODXL1, OTX2 or pluripotency markers (Supplementary information, Fig. S1f, g), consistent with the scRNA-seq data (Fig. 1e; Supplementary information, Fig. S1e). We also observed a few OCT4^−^KDR^+^ ExM cells without GATA6 expression (Supplementary information, Fig. S1f, g). Based on these data, we identified OCT4^−^SOX2^−^KDR^−^SOX17^+^FOXA1^+^PODXL1^+^ cells as XEN cells and OCT4^−^SOX2^−^SOX17^−^FOXA1^−^KDR^+^ as ExM cells.

We also performed KEGG pathway analysis to reveal signaling pathways involved in lineage developments. The AME, PS, ExM, VE/YE and AVE clusters are enriched with genes associated with BMP and WNT signaling (Fig. 1f), suggesting the important roles of BMP and WNT signaling in the post-implantation EPI and HB lineage specifications. Specifically, *BMP2* and *BMP4* are highly upregulated in the XEN and ExM, respectively, whereas BMP target genes *ID1*, *ID2*, *ID3* and *ID4* are mainly expressed in the AME and PS (Fig. 1f; Supplementary information, Fig. S1e), implying potential roles of XEN and ExM in inducing AME and PS specifications, which is further confirmed by intercellular communication analysis (Fig. 1i). *WNT6* is specifically expressed in AME, while *WNT3*, *WNT5A* and *WNT5B* are mainly expressed in PS (Fig. 1f; Supplementary information, Fig. S1e). Interaction analysis also showed that AME and PS are the major senders of WNT signaling (Supplementary information, Fig. S1h). Interestingly, WNT inhibitors *DKK1*, *SFRP1* and *SFRP2* are mainly upregulated in ExM, AVE and EPI, respectively (Supplementary information, Fig. S1e); however, *DKK1* and *SFRP1* are mainly expressed in monkey AVE and EPI, respectively ^28, 31^, suggesting different regulatory patterns of WNT signaling in humans and monkeys.

### Derivation of naive hESCs under normoxia condition

To model post-implantation HB and EPI lineage developments, we sought to establish an embryoid using naive hPSCs, which are the *in vitro* counterparts of the pre-implantation EPI (Pre-EPI). Naive hPSCs have been derived using t2iLGö, 5i/LAF or HENSM medium conditions under 5% O_2_ ^32–34^. Such a low oxygen level, however, causes pluripotency loss and cell death in human embryos cultured through the implantation stage ^1^. We thus first sought to derive naive hPSCs in normoxia (21% O_2_). In the previous works, we established the AIC medium, which is composed of modified N2B27 medium supplemented with AIC (Activin-A, IWP2 and CHIR99021), and supports efficient derivation and long-term expansion of primed hPSCs through single-cell passage ^35^. Naive hPSCs grown in the t2iLGö proliferate more rapidly and are karyotypically more stable than those cultured in 5i/LAF ^36–38^. We therefore singled out the key components of t2iLGö for the maintenance of naive pluripotency, including the MEK inhibitor PD0325901, the protein kinase C inhibitor Gö6983 and human LIF ^39^, which were incorporated into the AIC medium to establish a new culture system, termed AIC-N medium (Fig. 2a). AIC-N allows isolation and expansion of naive hESCs from blastocysts or their conversion from conventional primed hESCs in normoxia (Fig. 2b). AIC-N hESCs could be expanded for at least 50 passages in AIC-N and retained pluripotency properties similar to naive hPSCs established under 5% O_2_, in terms of their expression of pluripotency markers, transcriptome, methylation and potential to differentiate into embryonic and extraembryonic lineages (teratomas, blastoids and trophoblast lineages) (Fig. 2c-h; Supplementary information, Fig. S2a-h)^32–34, 40–42^.

**Fig. 2.**
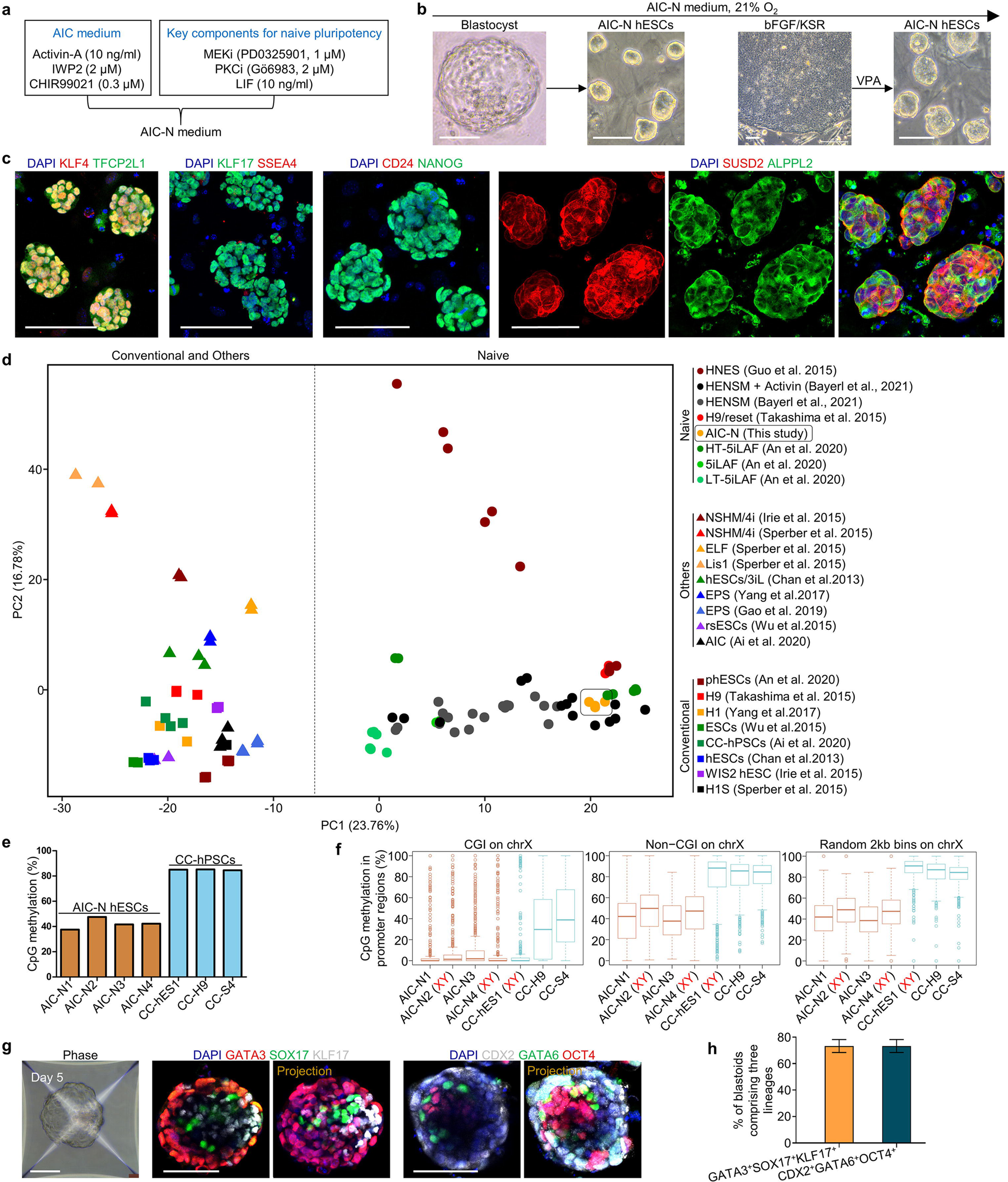
Derivation of naive hESCs under normoxia (21% O_2_). **a** Schematic of the AIC-N medium by incorporating the key components for the maintenance of naive pluripotency^39^ into the AIC medium^35^. **b** Representative contrast-phase images showing the generation of naive hESCs (AIC-N hESCs) from blastocysts and by conversion of primed hPSCs under normoxia. Four AIC-N hESC lines were established from seven blastocysts. The conversion of primed hPSCs required addition of valproic acid (VPA) for 4-6 days. **c** Immunostaining of naive, primed and general pluripotency markers for AIC-N hESCs. **d** PCA analysis of the gene expression profiles of hPSCs grown in various conditions. **e** Whole-genome CpG methylation levels of four AIC-N hESC lines and three primed hPSC (CC-hPSC) lines based on bisulfite sequencing (BS-seq) analysis. **f** CpG methylation levels at X-linked promoter CpG islands (CGIs) (left), non-CGI promoter regions (middle) and random 2 kb bins (right) in AIC-N hESCs and CC-hPSCs. The random 2kb bins do not overlap any CGIs or non-CG promoters. Promoters are defined as +/-1kb regions around transcription start sites. **g** Representative contrast-phase and immunostaining images showing the generation of blastoids from AIC-N hESCs. **h** Quantification of the percentage of blastoids comprising three lineages (trophectoderm-, hypoblast– and epiblast-like cells), three independent experiments; more than 100 blastoids were quantified for each experiment. Data are presented as mean±s.d. Scale bars, 100 µm. See also Supplementary information, Fig. S2.

### Generation and development of human E-assembloids

We next attempted to aggregate AIC-N hESCs with either human trophoblast stem cells (hTSCs) ^43^ or naive hESC-derived trophectoderm-like cells (nTEs) ^41, 42^ in a basal medium diluted with half hTSC medium (termed M1) supplemented with 1% Matrigel, for the development of an integrated embryo model. However, poor (∼3%) integrations of AIC-N hESCs with either hTSCs or nTEs were observed (Fig. 3a-c; Supplementary information, Fig. S3a-d). We then considered inducing trophoblast differentiation from primed AIC-hESCs ^35^ by sequential treatment of the cells with BMP4/SB431542 and the hTSC medium (Supplementary information, Fig. S3e) ^43^. The resulting cells, herein termed BMP4-induced cells (BICs), could be expanded over 10 passages and gradually acquire a hTSC-like identity over culture (Supplementary information, Fig. S3f, g). Gene expression patterns suggest that BICs undergo a fate transition to post-implantation trophoblast-like state over culture after exhibiting a transient amnion-like phenotype (Supplementary information, Fig. S3h-k). Interestingly, when assembling AIC-N hESCs and BICs different days after induction in M1 supplemented with 1% Matrigel, the wrapping capacity of BICs for AIC-N hESCs decreased with their extended culture, with D2 BICs showing the highest envelopment efficiency (over 90%) concomitant with efficient cell survival (Fig. 3a-c; Supplementary information, Fig. S3l).

**Fig. 3.**
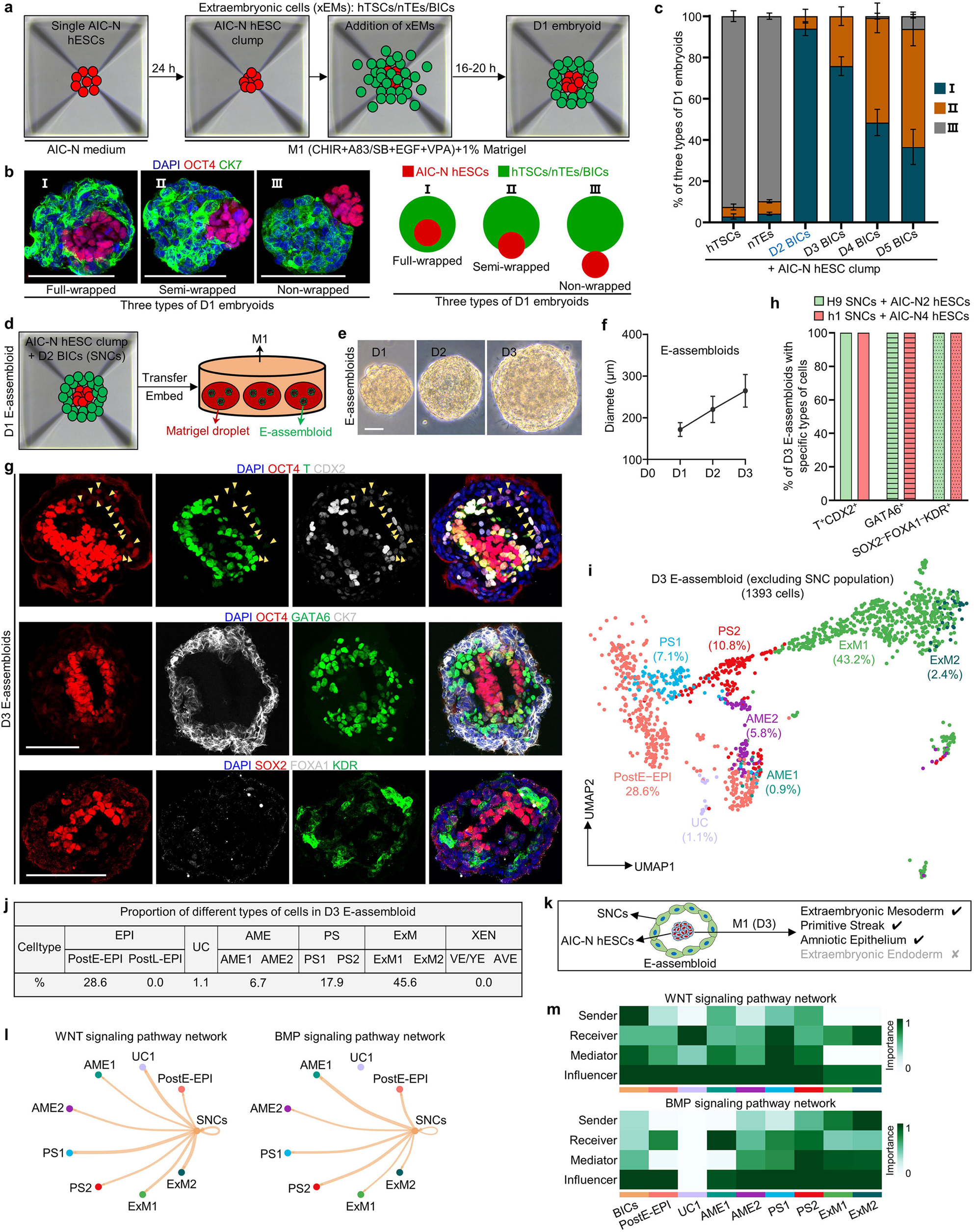
Generation and extended culture of E-assembloids. **a** Diagram for generating human embryoids by assembly of AIC-N hESCs and different types of extraembryonic cells (xEMs). hTSCs, human trophoblast stem cells; nTEs, naive hESC-derived trophectoderm-like cells; BICs, BMP4-induced cells. **b** Representative staining images (left) and schematics (right) depicting three types of D1 embryoids according to the state of AIC-hESCs wrapped by xEMs. CK7 for xEMs, and OCT4 for AIC-N hESCs. **c** Quantification of three types of D1 embryoids derived from AIC-N hESCs and different types of xEMs. Three independent experiments; more than 100 embryoids were quantified for each experimental group. Data are presented as mean±s.d. A large number of dead cells surround the embryoid assembled from D1 BICs (Supplementary information, Fig. S3l), therefore, which was not used for further study. **d** Diagram of extended culture of human E-assembloids assembled from AIC-N hESCs and D2 BICs (SNCs). **e** Representative contrast-phase images of E-assembloids during extended culture in the M1 condition. **f** Dynamic diameters of E-assembloids during extended culture in the M1 condition. N=3 independent experiments; at least 100 E-assembloids were quantified each experiment; data are presented as mean±s.d. **g** Immunostaining of D3 E-assembloids with specific lineage markers. Yellow arrowheads indicate AMELCs. **h** Quantification of different types of E-assembloids indicated in **g**. Three independent experiments; at least 100 E-assembloids were quantified each experiment; data are presented as mean±s.d. **i** UMAP visualization of integration analysis of the remaining cell types after excluding TrB and SNC population from human embryos and D3 E-assembloids, respectively. **j** Proportion of different subtypes of cells indicated in **i** based on analysis of scRNA-seq data. **k** Schematic showing the specifications of indicated cell lineages in D3 E-assembloids. **l** The inferred WNT and BMP signaling outputs from SNCs, line weight represents the communication probability. **m** Heatmap shows the relative importance of each cell group based on the computed four network centrality measures of WNT and BMP signaling network. EPI, epiblast; PostE−EPI and PostL−EPI, post-implantation early and late EPI; UC, undefined cell-type; AME, amniotic epithelium; PS, primitive streak; ExM, extraembryonic mesoderm; XEN, extraembryonic endoderm; VE/YE, visceral/yolk sac endoderm; AVE, anterior visceral endoderm; TrB, trophoblast; SNCs, signaling nest cells. Scale bars, 100 µm. See also Supplementary information, Fig. S3 and Fig. S4.

WNT and BMP signaling play key roles in early human embryonic development ^9, 14, 31, 44–46^. To investigate whether the extraembryonic tissues regulate the development of embryonic compartment by secreting WNT and BMP signals, we analyzed the WNT and BMP signaling interactions between extraembryonic and embryonic compartments in 3D-cultured human embryos before E10. As expected, the peripheral TrB and XEN send WNT and BMP signals to the internal EPI, respectively (Supplementary information, Fig. S4a, b). Similarly, D2 BICs, intermediates that express amnion markers transiently during trophoblast induction, show upregulated expression of WNT and BMP ligands (Supplementary information, Fig. S4c), suggesting that they may have a regulatory function of extraembryonic tissue for embryonic compartment. Together, even though it is unclear the counterpart of D2 BICs in the natural human embryo, they express key signaling ligand genes and provide an extraembryonic-like nest for AIC-N hESC clump, similar to the strategy reported in two recent studies using cells from hESCs induced by BMP4 to instruct epiblast development into early gastrulation cell types ^10^ or experimentally engineered morphogen signaling centre that functions as an organizer to instruct development of embryo-like entities (embryoids)^47^. We thus termed D2 BICs signaling nest cells (SNCs), and the embryo-like assembloid generated through integrating SNCs and naive hESCs were termed E-assembloid.

We next examined the developmental potential of E-assembloids. Under the M1 condition, E-assembloids embedded in Matrigel droplets gradually grew in size (Fig. 3d-f). On D3, a large number of T^+^/CDX2^+^ cells (putative primitive streak-like cells, PSLCs or amniotic epithelium-like cells, AMELCs), and GATA6^+^ ExM-like cells (ExMLCs) that expressed ExM markers KDR and COL6A1, but not XEN markers FOXA1, OTX2 and EOMES, emerged in E-assembloids (Fig. 3g, h; Supplementary information, Fig. S4d, e). The E-assembloids generated from different cell lines showed similar results (Fig. 3h). To confirm cell lineage origins, E-assembloids were generated using female SNCs and male AIC-N hESCs. Single-cell gene expression analysis of Y-chromosome specific gene *RPS4Y1* and lineage markers showed the presence of ExMLCs and PSLCs/AMELCs and their origins from AIC-N hESCs (Supplementary information, Fig. S4f), which was further confirmed by immunofluorescence staining of E-assembloids constructed from SNCs and mCherry-labeled AIC-N hESCs (Supplementary information, Fig. S4g). To characterize D3 E-assembloids, the scRNA-seq data from E10-14 human embryos were used as a reference for comparative transcriptome analysis. UMAP analyses revealed a clear distinction between SNC and TrB populations, which was further confirmed by differential expressions of trophoblast and amnion marker genes in the two populations (Supplementary information, Fig. S4h, i). Thus, SNCs in E-assembloids fail to recapitulate the development of TrB-related cell lineages. To determine how AIC-N hESC derivatives in E-assembloids correspond to their embryonic counterparts, we excluded the TrB and SNC populations from the scRNA-seq data of embryos and E-assembloids, respectively. Consistent with the immunostaining results, further integrated UMAP analyses showed that AIC-N hESC derivatives consisted of cells with molecular profiles comparable to PostE-EPI (28.6%), UC (1.1%), AME (6.7%), PS (17.9%) and ExM (45.6%), but lacked PostL-EPI-like cells and XEN-like cells (XENLCs) (Fig. 3i, j; Supplementary information, Fig. S4j). In the E-assembloids, there are no XENLCs, but a large number of ExMLCs (Fig. 3j), suggesting these ExMLCs as progenies of EPILCs, consistent with a recent study ^46^. Together, E-assembloids cultured in the M1 condition recapitulate the development of human ExM, PS and AME but not PostL-EPI and XEN (Fig. 3k).

### WNT and BMP signaling orchestrate post-implantation lineage development

In contrast to E-assembloids, most of AIC-N hESC clumps grown alone in the M1 condition maintained pluripotency even on D9 (Supplementary information, Fig. S4k, l), suggesting that there are inductive signals from SNCs to drive E-assembloid development. Since SNCs express WNT and BMP ligands (Supplementary information, Fig. S4c), and the specifications of ExM, PS and amnion are closely related to WNT and BMP signaling ^8–10, 14, 46^. To investigate whether the SNCs provide WNT and BMP signals to induce embryonic and extraembryonic lineage differentiation in E-assembloids, we analyzed the intercellular communication networks in D3 E-assembloids by single-cell transcriptomes, and confirmed that BICs do produce WNT and BMP signals that are received by epiblast-like cells (EPILCs) and other types of cells (Fig. 3l, m). We therefore manipulated WNT and BMP signaling to assess their effects on the developments of E-assembloids, AIC-N hESC clumps and human embryos. Specifically, we first inhibited WNT and BMP pathways using chemical inhibitors during extended culture of E-assembloids (Fig. 4a). Contrary to the development of E-assembloids in the M1 condition, inhibition of either WNT or BMP signaling in E-assembloids largely delayed EPILC development toward embryonic and extraembryonic lineages, specifications of which were even completely blocked by dual inhibitions of WNT and BMP signaling (Fig. 4a, b; Supplementary information, Fig. S5a, b). Notably, both WNT and BMP inhibitions significantly reduced ExM specification, but with differential effects on AME and PS specifications (Fig. 4a-d; Supplementary information, Fig. S5a, c, d). WNT inhibition abolished PS specification but promoted AME development. In contrast, BMP inhibition abolished AME development and reduced PS specification (Fig. 4a-d; Supplementary information, Fig. S5a, c, d). Consistent with chemical inhibition assays, developments of ExM, AME and PS were significantly inhibited in E-assembloids generated from BMP4^-/-^/7^-/-^ SNCs (Fig. 4e, f; Supplementary information, Fig. S5e, f). To further validate the functions of WNT and BMP signals, different combinations of WNT and BMP agonists and inhibitors were used to treat AIC-N hESC clumps that were not wrapped in SNCs (Fig. 4g). The results showed that ExM specification required dual activation of WNT and BMP signaling, and PS and AME specifications required activation of WNT and BMP signaling, respectively (Fig. 4h; Supplementary information, Fig. S5g). Finally, we used 3D-cultured human embryos ^4^ to further confirm that dual activation of WNT and BMP signals is indispensable for ExM specification (Supplementary information, Fig. S5h). Together, our results demonstrated that ExM specification required both WNT and BMP signaling, whereas WNT or BMP signaling is required for PS or AME development, respectively (Fig. 4i).

**Fig. 4.**
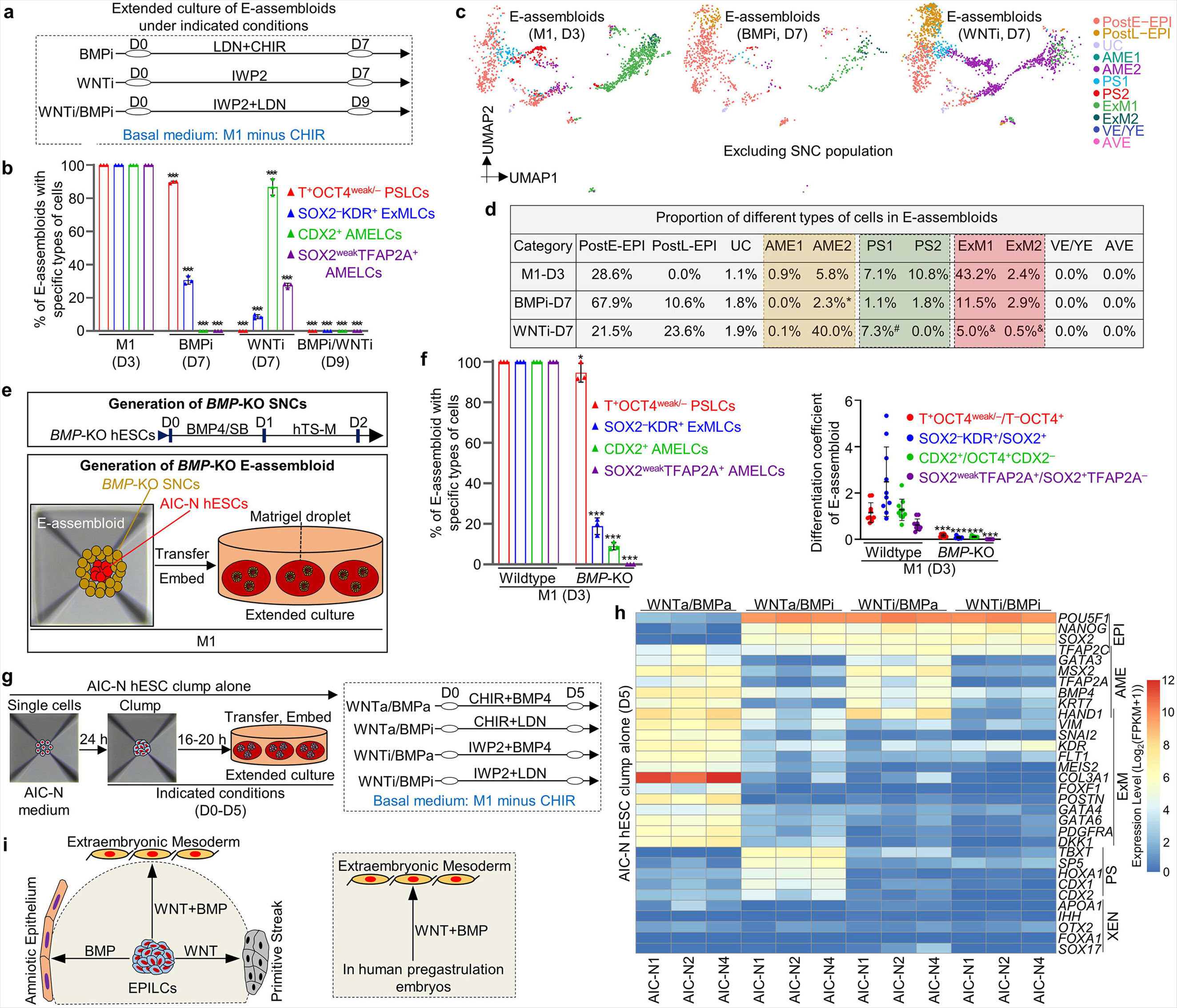
WNT and BMP signaling orchestrate lineage development. **a** Schematic diagram of different culture conditions for E-assembloids. BMPi and WNTi represent BMP inhibition and WNT inhibition, respectively. Compared to the M1 condition, BMP or/and WNT inhibition delayed/blocked EPILC development in E-assembloids toward embryonic and extraembryonic lineages, we therefore prolonged culture of E-assembloids to observe the potential effects of WNT or/and BMP inhibition (seven days for individual inhibition and nine days for dual inhibition). **b** Quantification of different types of E-assembloids cultured in indicated conditions at indicated time points. N=3 independent experiments; at least 100 E-assembloids were quantified each experiment; data are presented as mean±s.d. *** p≤0.001 with Student’s *t*-test. **c** UMAP visualization of integration analysis of the remaining cell types after excluding TrB and SNC population from human embryos and E-assembloids grown in indicated conditions, respectively. **d** Proportion of different cell subtypes in E-assembloids cultured in indicated conditions at different time points, see also Supplementary information, Fig. S5d. * These cells were clustered into AME2, but weakly co-expressed AME, PS and ExM marker genes, indicating a uncertain identity; ^#^ these cells were clustered into PS1 and co-expressed pluripotency and AME but not PS marker genes, implying a nascent amnion identity; ^&^ these cells were clustered into ExM1/2 and highly expressed AME but not ExM marker genes, implying a amnion identity. **e** Diagram of generation of *BMP*-KO H9 SNCs (top) and *BMP*-KO E-assembloids (bottom), see also Supplementary information, Fig. S5e. **f** Porportion (left) and differentiation coefficient (right) of different types of E-assembloids (wildtype and *BMP*-KO) grown in the M1 condition on D3. The differentiation coefficient represents the ratio of differentiated cells to pluripotent cells. For the porportion, n=3 independent experiments; at least 100 E-assembloids were quantified each experiment; for the differentiation coefficient, 10 E-assembloids were randomly selected for statistics in each group. Data are presented as mean±s.d. * p≤0.05 and *** p≤0.001 with Student’s *t*-test. **g** Schematic diagram of AIC-N hESC clumps cultured alone in different culture conditions for five days. WNTa and BMPa represent WNT activation and BMP activation, respectively. **h** Heatmap of representative marker genes of different lineages from three AIC-N hESC lines cultured in indicated conditions. **i** Functional diagram of WNT and BMP signaling on embryonic lineage development. CHIR, CHIR99021; LDN, LDN193189-2HCl; SB, SB431542; EPI, epiblast; PostE−EPI and PostL−EPI, post-implantation early and late EPI; UC, undefined cell-type; AME, amniotic epithelium; PS, primitive streak; XEN, extraembryonic endoderm; ExM, extraembryonic mesoderm; VE/YE, visceral/yolk sac endoderm; AVE, anterior visceral endoderm; TrB, trophoblast; SNCs, signaling nest cells. Scale bars, 100 µm. See also Supplementary information, Fig. S5.

### Signaling regulatory mechanisms controlling extraembryonic endoderm specification

Even with modulated WNT and BMP signaling and extended culture, E-assembloids and AIC-N hESC clumps did not give rise to XENLCs (Fig. 4d, f, h). Since Nodal signaling is activated in XEN (Fig. 5a, b), we asked whether the presence of Activin/Nodal inhibitors (A83-01+SB431542, AS) in the M1 medium might have blocked XENLC specification in E-assembloids and AIC-N hESC clumps. To this end, we first induced 2D differentiation of AIC-N hESCs with or without AS (Fig. 5c), with results showing that CHIR+BMP4 induced differentiation of AIC-N hESCs into a heterogeneous population containing both ExMLCs and XENLCs (Fig. 5d, Conditions (2) and (3)). Importantly, inhibition of endogenous Nodal signaling with AS blocked XENLC specification (Fig. 5d, Conditions (1) and (4)). Interestingly, the effect of CHIR+BMP4 treatment was time-dependent: a transient (2 days) induction of CHIR+BMP4 was beneficial for XENLC specification (Fig. 5d, e, Condition (3)), whereas their continuous (4 days) treatment promoted ExMLC differentiation at expense of XENLCs (Fig. 5d, Condition (2)). Notably, activation of WNT or BMP signaling alone was ineffective in specifications of XENLCs and ExMLCs (Fig. 5f), suggesting that their specifications require a synergy between WNT and BMP signaling. Together, effective differentiation of AIC-N hESCs into XENLCs requires short-time activation of both WNT and BMP signaling in a Nodal-dependent manner (Fig. 5g, h).

**Fig. 5.**
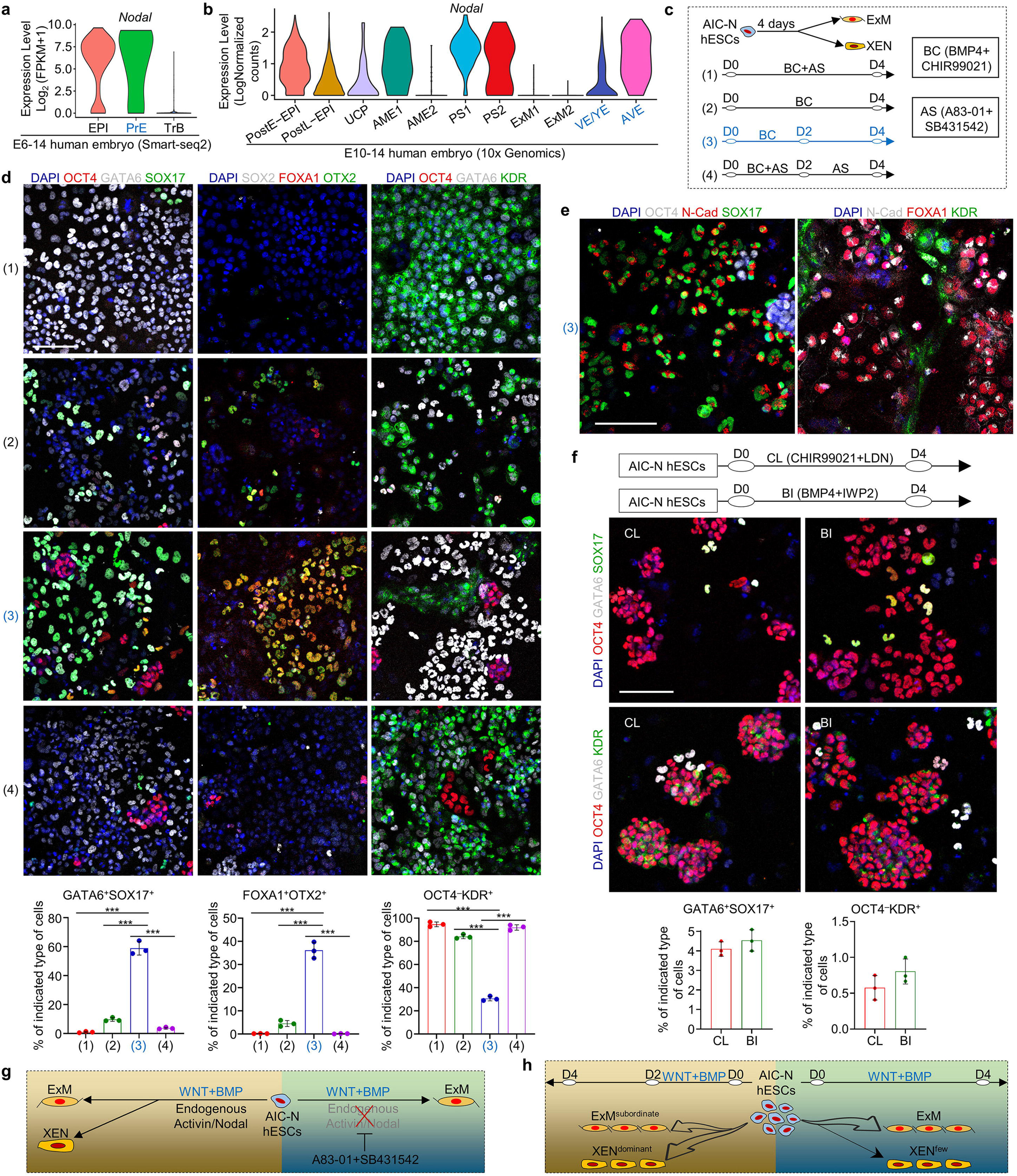
Extraembryonic endoderm specification. **a** and **b** Violin plots of Nodal gene expressed in different types of cells by scRNA-seq data. **a** published smart-seq2 data ^4^, **b** 10x Genomics data. **c** Functional experiment of WNT, BMP and Nodal signals on XENLC and ExMLC specifications. **d** Representative staining images showing the specifications of XENLCs and ExMLCs and proportion of indicated cell type under different culture conditions. N=3 independent experiments; data are presented as mean±s.d. *** p≤0.001 with Student’s t-test. **e** Representative staining images showing the specifications of XENLCs and ExMLCs under Condition (3). **f** Schematic of culture conditions (top), representative staining images (middle) and proportion of different cell type (bottom) under indicated culture conditions. N=3 independent experiments; data are presented as mean±s.d. **g** and **h** Functional schematic diagram of WNT, BMP and Nodal signals on the specifications of XENLCs and ExMLCs. EPI, epiblast; PrE, primitive endoderm; TrB, trophoblast; PostE−EPI and PostL−EPI, post-implantation early and late EPI; UC, undefined cell-type; AME, amniotic epithelium; PS, primitive streak; ExM, extraembryonic mesoderm; VE/YE, visceral/yolk sac endoderm; AVE, anterior visceral endoderm; XEN, extraembryonic endoderm; LCs, –like cells; Scale bars, 100 µm.

### Establishment of an improved E-assembloid by regulating signaling pathways

Based on the principle of specifications of embryonic and extraembryonic lineages involving WNT, BMP and Nodal signaling (Fig. 4i and 5g, h), we established an improved Protocol for assembly and extended culture of E-assembloids (Fig. 6a). The improved Protocol was based on the M1 minus Activin/Nodal inhibitors and included three steps: (1) lineage induction by endogenous signals secreted from SNCs and exogenous WNT agonist (1 μM CHIR) from D0 to D2; (2) inhibition of excessive ExM specification by blocking endogenous BMP signals with LDN and removing exogenous WNT agonist (CHIR withdrawal) from D2 to D4; and (3) spontaneous lineage specification and self-organization without exogenous signal interference from D4 onwards (Fig. 6a). This improved Protocol allowed E-assembloids to progressively grow and develop with similar sizes to 3D-cultured human embryos (Fig. 6b; Supplementary information, Fig. S6a). Remarkably, immunostaining for lineage markers revealed dynamic development of XENLCs and ExMLCs in E-assembloids (Fig. 6c, d; Supplementary information, Fig. S6b-d). On D1, no XENLCs or ExMLCs were observed (Fig. 6d; Supplementary information, Fig. S6b). On D2, abundant GATA6^+^OCT4^weak^NANOG^weak^ cells and small numbers of SOX17^+^BLIMP1^+^ cells were detected in over 90% of E-assembloids; few FOXA1^+^, OTX2^+^, SOX2^−^FOXA1^−^KDR^+^ and OCT4^weak^ KDR^+^ cells also were specified in approximately 10%, 20%, 10% and 10% E-assembloids, respectively; and no PODXL1^+^GATA6^+^ XENLCs and BST2^+^ ExMLCs were observed (Fig. 6d; Supplementary information, Fig. S6b-d), implying an initial XENLC and ExMLC specifications. On D3, more than 90% of E-assembloids produced abundant SOX17^+^OTX2^+^ and GATA6^+^PODXL1^+^ XENLCs and a few OCT4^weak/–^KDR^+^ ExMLCs, and approximately 80% of E-assembloids gave rise to small numbers of BST2^+^ ExMLCs. On D4, almost all E-assembloids contained abundant GATA6^+^, FOXA1^+^, SOX17^+^BLIMP1^+^, PODXL1^+^GATA6^+^ XENLCs that surrounded OCT4^+^ EPILC compartment, accompanied by specification of SOX2^−^KDR^+^ ExMLCs (Fig. 6d; Supplementary information, Fig. S6b). Together, during XENLC and ExMLC specifications in E-assembloids, GATA6 is first activated; followed by SOX17 and BLIMP1; and OTX2, FOXA1, KDR, PODXL1 and BST2 are activated last.

**Fig. 6.**
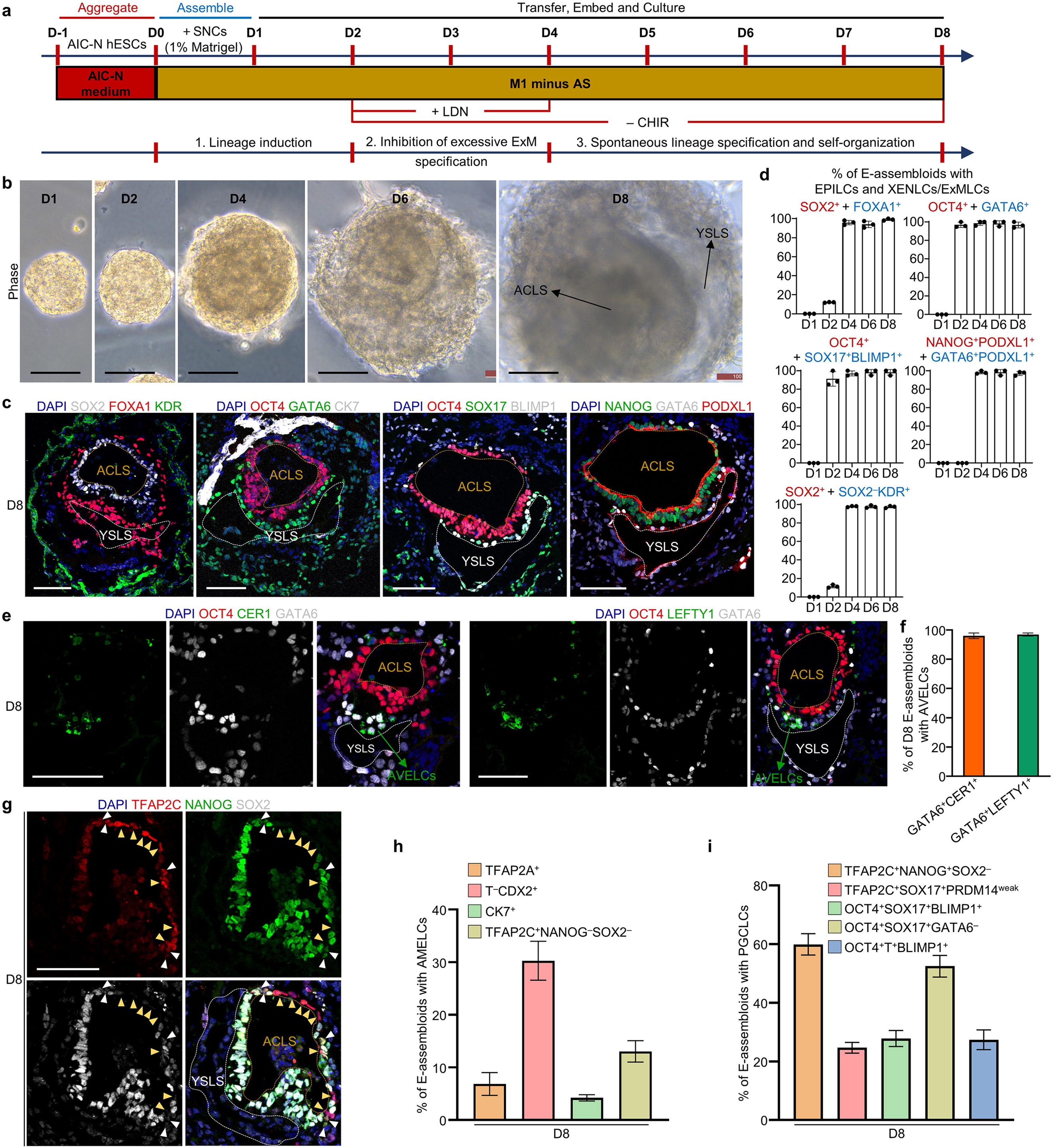
E-assembloids recapitulate early post-implantation embryogenesis. **a** Schematic representation of improved Protocol for assembly and extended culture of E-assembloids. AS, A83-01+SB431542; CHIR, CHIR99021; LDN, LDN193189-2HCl; ‘+’ and ‘–’ represent ‘add’ and ‘withdraw’, respectively. **b** Representative contrast-phase images of E-assembloids during extended culture. **c** Representative immunostaining images showing the formation of ACLS and YSLS in D8 E-assembloids. **d** Quantification of the E-assembloids with EPILCs (red) and XENLCs/ExMLCs (blue) at different time points. N = 3 independent experiments; more than 100 structures were quantified each experiment; data are presented as mean±s.d. **e** Representative immunostaining images showing the generation of AVELCs in D8 E-assembloids. **f** Quantification of the E-assembloids with AVELCs indicated in (E). N = 3 independent experiments; more than 100 structures were quantified each experiment; data are presented as mean±s.d. **g** Representative immunostaining images showing the specifications of AMELCs (yellow arrowheads) and PGCLCs (white arrowheads) in D8 E-assembloids. **h** and **i** Quantification of D8 E-assembloids containing AMELCs and PGCLCs. N = 3 independent experiments; more than 100 structures were quantified each experiment; data are presented as mean±s.d. EPI, epiblast; XEN, extraembryonic endoderm; ExM, extraembryonic mesoderm; AVE, anterior visceral endoderm; AME, amniotic epithelium; PGC, primordial germ cell; AC, amniotic cavity; YS, yolk sac; LS, –like structure; LCs, –like cells. Scale bars, 100 µm. See also Supplementary information, Fig. S6.

On D8, > 90% of E-assembloids contained two expanded cavities, the yolk sac-like and amniotic cavity-like structures, surrounded by ExMLCs (Fig. 6b-d). In > 90% of E-assembloids, notably, we observed XENLCs expressing CER1 and LEFTY1 (Fig. 6e, f), suggestive of an AVE-like identity ^24, 31^. Similar results were obtained from the E-assembloids constructed from two different cell lines (Supplementary information, Fig. S6e). These results demonstrated that the XENLC deficiency in E-assembloids was solved by the improved Protocol (Fig. 6a). Furthermore, AMELCs were specified at the prospective dorsal pole of the amniotic cavity-like structure, and PGCLCs were detected within AME-like tissue or near the junction of EPILCs and XENLCs (Fig. 6g-i; Supplementary information, Fig. S6f, g). In addition, in approximately 20% of E-assembloids, some T^+^ putative PSLCs are observed (Supplementary information, Fig. S6h, i). However, in addition to being at the putative posterior junction of the yolk sac-like structure and EPILC compartment, some of these putative PSLCs, which co-expressed CDX2, also appeared on opposite side of the putative posterior domain (Supplementary information, Fig. S6f), exhibiting a disorganized localization. To answer whether the effects of BICs can be simulated by adding exogenous agonists of WNT and BMP signaling, we transiently treated AIC-NhESC clumps with CHIR+BMP4 for 12 hours (Supplementary information, Fig. S6j), and resulted in the generation of a large number of embryonic and extraembryonic lineages on D6, including EPILCs (OCT4^+^NANOG^+^SOX2^+^), PSLCs and AMELCs (T^+^CDX2^+^ and TFAP2A^+^), XENLCs (GATA6^+^FOXA1^+^) and ExMLCs (GATA6^+^FOXA1^−^ and SOX2^−^KDR^+^) (Supplementary information, Fig. S6k, l), showing exogenous activation of WNT and BMP signaling could partially simulate the induction effect of BICs. However, in these resulting AIC-N hESC spheres, the different types of cells are mostly arranged irregularly (Supplementary information, Fig. S6k). Therefore BICs, in addition to their effect of inducing different lineage specifications, may also provide mechanical effects that favor ordered organization of different cell lineages.

To further characterize D8 E-assembloids, comparative transcriptome analysis of integrated scRNA-seq data of 3D-cultured human embryos and D8 E-assembloids was conducted (Supplementary information, Fig. S7a). UMAP plots and differential expressions of trophoblast and amnion marker genes reveal an obvious difference between SNC and TrB populations (Supplementary information, Fig. S7a, b), but similar compositions of HB– and EPI-related lineages in human embryos and E-assembloids, including postE-EPI, postL-EPI, UC, AME2, PS1/2, ExM1/2, VE/YE and AVE (Fig. 7a; Supplementary information, Fig. S7c, d). AME1, however, was not detected by scRNA-seq in E-assembloids (Fig. 7a). We further split the integrated UMAP data of 3D-cultured human embryos according to different developmental time points and found that the D8 E-assembloids are similar to the E13/14 embryos in terms of cell lineage compositions (Supplementary information, Fig. S7e). Although *AFP*, *TTR*, *APOA4*, *APOC3* and *MTTP* were only expressed in a small number of VE/YE-like cells (VE/YELCs) in E-assembloids, very similar expression patterns of lineages and signaling regulatory genes were observed in other populations between human embryos and E-assembloids (Fig. 7b, c; Supplementary information, Fig. S7f). For BMP and WNT signal ligands, *BMP2* was mainly expressed in VE/YELCs and AVELCs; *BMP4* in ExMLCs, PSLCs and AMELCs; *WNT5A* in ExM1LCs and PS2LCs; *WNT5B* in PSLCs; and *WNT6* in AME2LCs (Fig. 7c). Intercellular communication analysis further showed that ExMLCs, AMELCs, XENLCs and PSLCs are the main senders of BMP signaling, while AMELCs and PSLCs are also the main senders of WNT signaling (Supplementary information, Fig. S7g). For antagonists of WNT, BMP and Nodal signalling, *HHEX*, *CER1*, *NOG* and *LEFTY1/2* were mainly expressed in AVELCs; *SFRP1* in AVELCs; and *SFRP2* in EPILCs (Fig. 7c). These results suggest that extensive intercellular interactions between different cell lineages exist in E-assembloids, similar to the human embryos (Fig. 1i; Supplementary information, Fig. S1h and S7g). Interestingly, in the ExM2/ExM2LC populations, we observed a population of special cells, which were negative for *POSTN*, *BMP4* and *BMP2*, but highly expressed *SOX17*, *HHEX* and haemato-endothelial markers *KDR*, *S100A6*, *PECAM1*, *MEF2C*, *CD34* and *CDH5* (Fig. 7d; Supplementary information, Fig. S7h), possibly suggesting a initial hematopoiesis in both human embryos and E-assembloids.

**Fig. 7.**
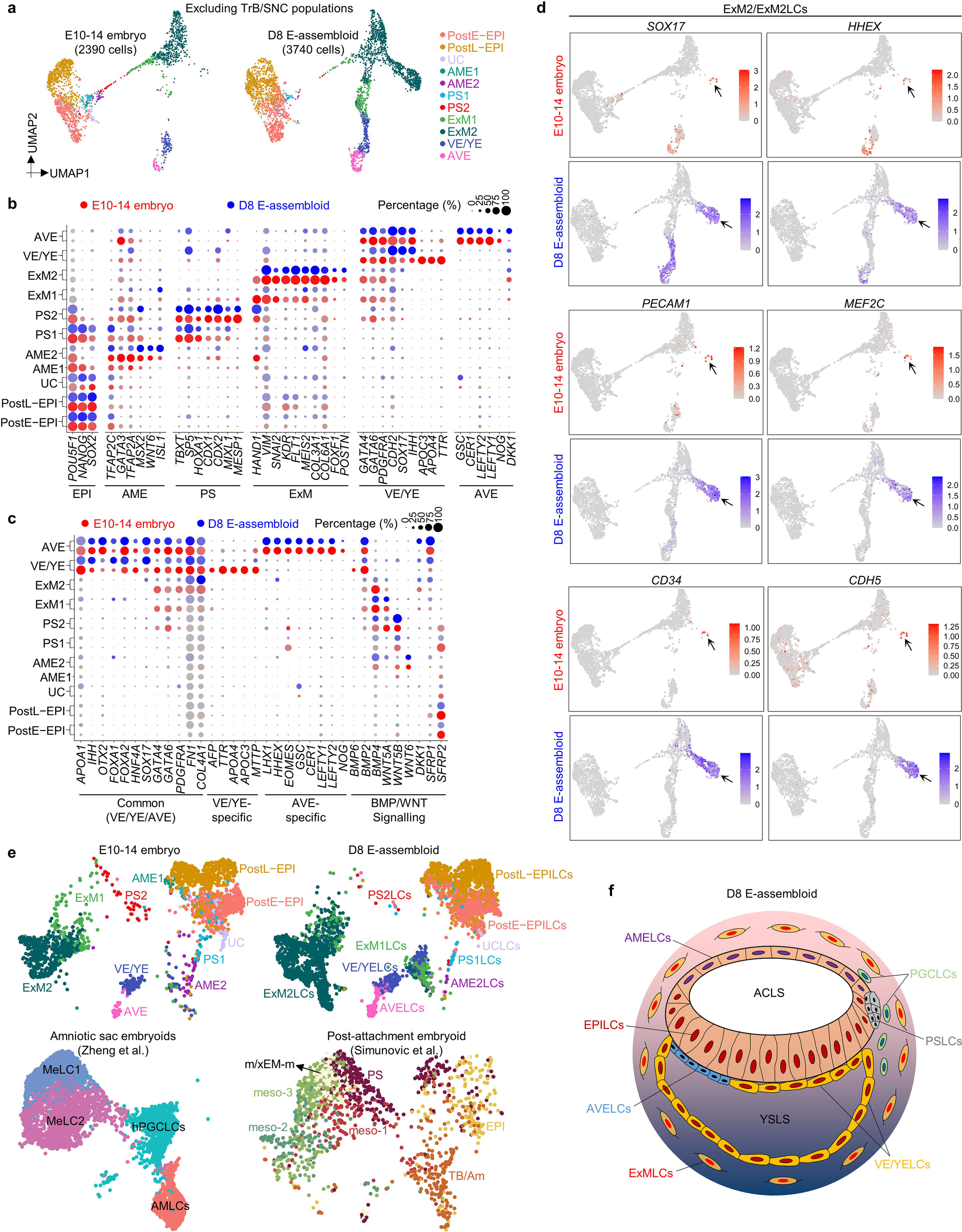
Comparative single-cell transcriptomics of E-assembloids and 3D-cultured embryos. **a** UMAP visualization of integration analysis of the remaining cell types after excluding TrB and SNC population in human embryos and D8 E-assembloids, respectively. **b** and **c** Dot plots of candidate genes specific for indicated cell subtypes in human embryo and D8 E-assembloid. **d** UMAP plots of indicated genes expressed in D8 E-assembloids and 3D-cultured embryos. Some cells in ExM2 population exhibit haemato-endothelial characteristics (arrow). **e** Comparative integration analysis of scRNA-seq datasets from 3D-cultured human embryos, E-assembloids and two other embryo models ^10, 14^. **f** Schematics showing 3D morphological feature of D8 E-assembloid. EPI, epiblast; PostE−EPI and PostL−EPI, post-implantation early and late EPI; UC, undefined cell-type; AME, amniotic epithelium; PS, primitive streak; ExM, extraembryonic mesoderm; VE/YE, visceral/yolk sac endoderm; AVE, anterior visceral endoderm; TrB, trophoblast; SNC, signaling nest cell; hPGCLCs, human primordial germ cell-like cells; MeLC, mesoderm-like cell; AMLCs, amniotic ectoderm-like cells; m/xEM-m, mesoderm/extraembryonic mesoderm; meso, mesoderm; TB/Am, trophoblast/amnion; LCs, –like cells. See also Supplementary information, Fig. S7.

## Discussion

In this study, we provided a complete cell atlas of peri-implantation embryonic lineages and identified XEN and ExM markers. Our data can serve as a resource useful for benchmarking human embryo models and embryonic/extraembryonic lineages *in vitro*. The bi-laminar disc, amniotic cavity, yolk sac and outer ExM are vital tissue structures in human pre-gastrulation embryos, but how these tissues develop together remains unclear. Amniotic sac embryoids developed by Zheng et al. reconstructe the formation of amniotic cavity or bipolar embryonic sac, and the specifications of early gastrulating cells and hPGCLCs, but cannot recapitulate the development of yolk sac and ExM (Fig. 7e) ^14^. Although the recently reported post-attachment embryoid (also known as embryo assembloid) provide a novel strategy for studying the anteroposterior symmetry breaking and onset of gastrulation in the human epiblast, this model cannot mimic the formation and specification of amniotic cavity, yolk sac and ExM (Fig. 7e)^10^. Our E-assembloid, a model of peri-implantation human development, can recapitulate developmental hallmarks and 3D structures of human pre-gastrulation EPI and HB derivatives, especially the bilaminar embryonic disc consisting of the amniotic cavity and the yolk sac, and the surrounding ExMs, which have never been presented simultaneously in previous embryo models (Fig. 7e, f). These advanced features place our E-assembloid as an important model for decoding developmental mechanisms of human peri-implantation embryogenesis, such as the interactions between embryonic and extraembryonic tissues, pluripotent state transitions in the epiblast ^48^, XEN specification, BMP signal source that drives amnion specification, and functions of specific genes and signaling pathways. Human ExM is specified prior to gastrulation and its origin remains currently unclear ^49, 50^. In 3D-cultured human embryos, RNA velocity map appeared to support EPI or early PS origin of ExM (Fig. 1g), but clustering showed that ExM is more closely related to XEN (Fig. 1h). Our results and a recently reported study have shown that EPILCs can differentiate into ExMLCs without XENLC specification (Fig. 3k, 4i, 5g) ^46^; however, whether ExM can originate from XEN is still unknown. With abundant XENLCs and ExMLCs (Fig. 7f), combined with live imaging and lineage tracing, our E-assembloid will provide an important platform for deciphering the developmental origin of ExM. Due to the low efficiency of trophoblast-related cells in wrapping naive hESCs, SNCs were used to construct E-assembloid. Thus, E-assembloid is not suitable for modeling peri-implantation human trophoblast development. Compared to complete human embryo models such as blastoids ^17–23^, a lack of trophoblast-related cell lineages in E-assembloids makes the model less ethically concerning. Therefore, whether the E-assembloid can continue to develop and mimic human gastrulation and organogenesis is well worth further exploring.

Using 3D-cultured human embryos and E-assembloids, as well as 2D and 3D differentiation of AIC-N hESCs, we systematically dissected the functions of WNT, BMP, and Nodal signaling on the specifications of PS, AME, ExM and XEN, and deciphered the mechanisms underlying XEN and ExM specifications (Fig. 4i, 5g, h). Our findings will be useful for improving differentiation protocols of naive hPSCs into XENLCs ^44^ or ExMLCs ^46^. In contrast to ExMLCs, our data show that specification of XENLCs functionally requires endogenous Nodal signaling, consistently with high *Nodal* expression in human XEN (Fig. 5a, b) ^51^. Although it has been reported that Nodal inhibition does not affect the number of GATA6^+^ cells in human and marmoset preimplantation embryos ^45, 52^, another study has shown that SOX17^+^ hypoblast cells are undetectable in Nodal inhibitor-treated human blastocysts ^53^. These seemingly conflicting results may be caused by different choices of markers – SOX17 marks hypoblast/XEN more specifically than GATA6 ^54^.

Our study has several limitations. First, the mechanisms affecting the capacity of extraembryonic cells (xEMs) to wrap naive hESC clumps are not revealed. A previous study proposed that robust patterning during tissue morphogenesis results from cellular self-organization based on a combinatorial adhesion code instructed by morphogen-directed gradient ^55^. The view is in line with the hypothesis about the cell sorting model of differential adhesion proposed by Steinberg in 1970 ^56^ and has recently been further validated in mouse synthetic embryos ^57^. Therefore, detecting differential adhesion molecules between naive hESCs and xEMs may be beneficial to answer why SNCs, rather than hTSCs and nTEs, efficiently wrap naive hESC clumps. Second, although SNCs can gradually acquire a hTSC-like identity in hTSC medium ^43^ under 2D culture condition, they cannot effectively give rise to TrB-related lineages in the extended cultured E-assembloids and therefore fail to recapitulate TrB development in human embryos. The results show that the fate of SNCs is influenced by the culture conditions and growing environment, but the underlying molecular mechanisms are unknown. Finally, in the extended cultured E-assembloid, it is not fully understood how the endogenous signals secreted by SNCs and exogenous signal interference truly represent signaling events during human peri-implantation development. These limitations are largely due to the lack of *in vitro* counterparts of human trophectoderm (TE) and HB. The establishment of faithful *in vitro* counterparts of TE and HB, combined with the strategy of mouse ETX embryo approach ^58^, may help overcome the current challenges in developing complete human embryo model.

In conclusion, this study offers important information for advancing knowledge of human embryology and reproduction, and the E-assembloid provides a useful model for future exploration of lineage diversification, signalling interactions and tissue patterning during human peri-implantation development.

## Acknowledgements

This work was supported by the National Key Research and Development Program of China (2022YFA1103100 and 2021YFA0805700), the National Natural Science Foundation of China (32130034, 82192874 and 82160294), the National Key Research and Development Program of China (2020YFA0112700), Major Basic Research Project of Science and Technology of Yunnan (202102AA100007, 202001BC070001 and 2019FY002), the Joint Special Funds for the Department of Science and Technology of Yunnan Province□Kunming Medical University (202101AY070001-260 and 202301AY070001-298). We thank Yan Bi for sharing ALPPL2 antibody. We acknowledge Ran Yang for details of the single cell dissociation and preparation of human embryos.

## Author contributions

T.L. and W.J. initiated the project. T.L. supervised the entire project. T.L. and Z.A. designed the experiments and wrote the manuscript. T.L. supervised Z.A., Y.Y., K.D. and B.N. to perform data analysis. Z.A., B.N., Y.Y., S.W., Y.H., C.Z., C.Z., R.K., T.C. W.L., N.L. and S.Z. performed stem cell isolation and culture, embryoid assembly and identification, culture system development and mechanistic studies. L.X., G.S., L.R., X.M., Y.L. and Z.W. performed human embryo culture. Y.Y., Z.A., K.D., Y.G. and W.L. analyzed sequencing data. Y.Z. and J.F. contributed to sequencing data analysis and manuscript preparation.

## Competing interests

The authors declare no competing interests.

---

**Fig. S1.**
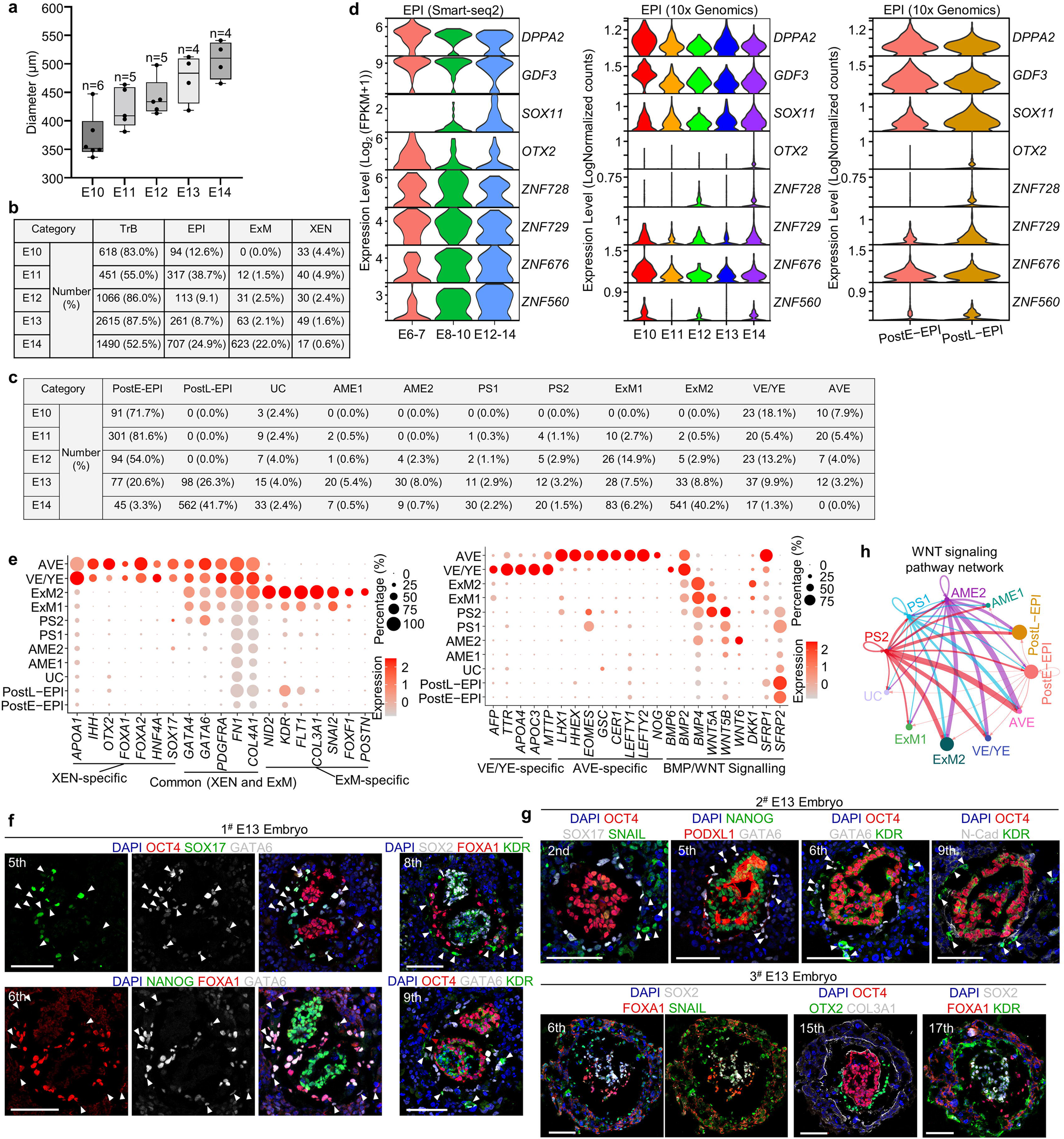
Single-cell transcriptomic profiling of 3D-cultured human post-implantation embryos, related to Fig. 1. **a** Diameters of 3D-cultured human embryos used for 10x Genomics sequencing. From embryonic day (E10-14), the number of 3D-cultured human embryos (corresponding thawed blastocysts) collected at each time point was 6 (10), 5 (10), 5 (12), 4 (14) and 4 (17) at each time point, respectively. **b** and **c** Numbers and proportions of different subtypes of cells in extended cultured human embryos at different developmental time points according to the results of scRNA-seq data. **d** Violin plots of indicated genes expressed in EPI by scRNA-seq data. The smart-seq2 results are from published data ^4^. **e** Dot plots of candidate genes specific for different cell subtypes. **f** and **g** Immunostaining of EPI, XEN and ExM markers in three E13 extended cultured human embryos. White ordinal numbers indicate section numbers, arrowheads indicate ExM cells, and red arrowheads indicate few OCT4^−^GATA6^−^KDR^+^ ExM cells. **f** a twin embyo; **g** two singleton embryos. We cultured ten E6 blastocysts and obtained three normally developed E13 embryos for immunostaining. **h** The inferred WNT signaling pathway network. Circle sizes are proportional to the number of cells in each subpopulation and line weight represents the communication probability. E, embryonic day; EPI, epiblast; PostE−EPI and PostL−EPI, post-implantation early and late EPI; UC, undefined cell-type; AME, amniotic epithelium; PS, primitive streak; ExM, extraembryonic mesoderm; VE/YE, visceral/yolk sac endoderm; AVE, anterior visceral endoderm; XEN, extraembryonic endoderm. Scale bars, 100 µm.

**Fig. S2.**
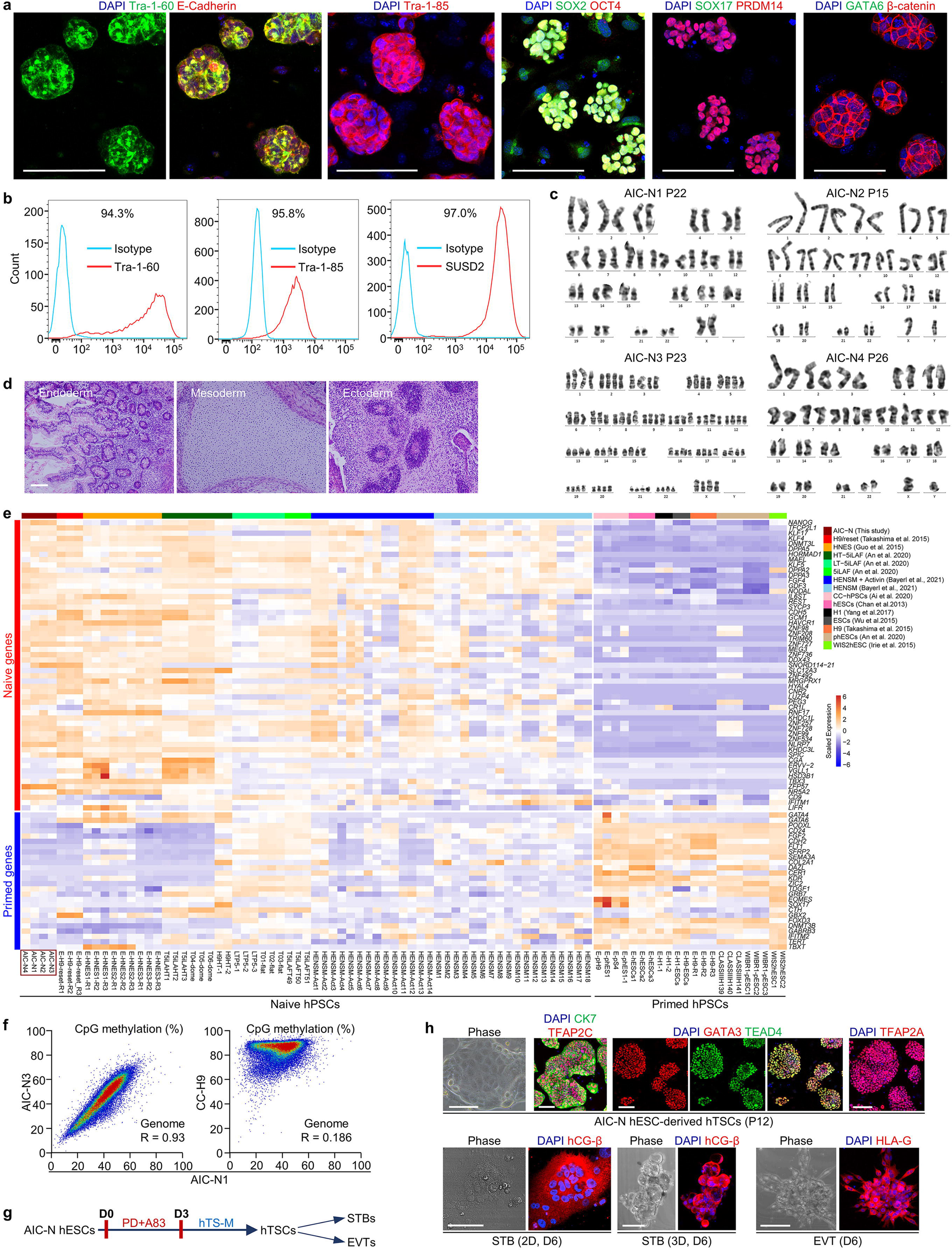
Identification of naïve hESCs, related to Fig. 2. **a** Immunostaining of general pluripotency and hypoblast markers for AIC-N hESCs. **b** Flow cytometry analysis of Tra-1-60, Tra-1-85 and SUSD2 in AIC-N hESCs. **c** G-banding karyotype analysis of four different AIC-N hESC lines. The passage (P) number for karyotyping is indicated. **d** AIC-N hESCs gave rise to teratomas including three germ lineages. **e** Heatmap of representative primed and naïve genes in hPSCs cultured in different conditions. Values represent log_2_ (FPKM+1) scaled by gene expression across samples. **f** Comparisons of global methylation in AIC-N1 (female), AIC-N3 (female) and CC-H9 (female, conventional) hESCs by averaging CpG methylation levels over 50-kb windows. **g** Differentiation schematic of trophoblast lineages from AIC-hESCs. **h** Representative contrast-phase and immunostaining images of different types of trophoblast cells differentiated from AIC-N hESCs. STB, syncytiotrophoblast-like cells; EVT, extravillous cytotrophoblast-like cells. Scale bars, 100 µm.

**Fig. S3.**
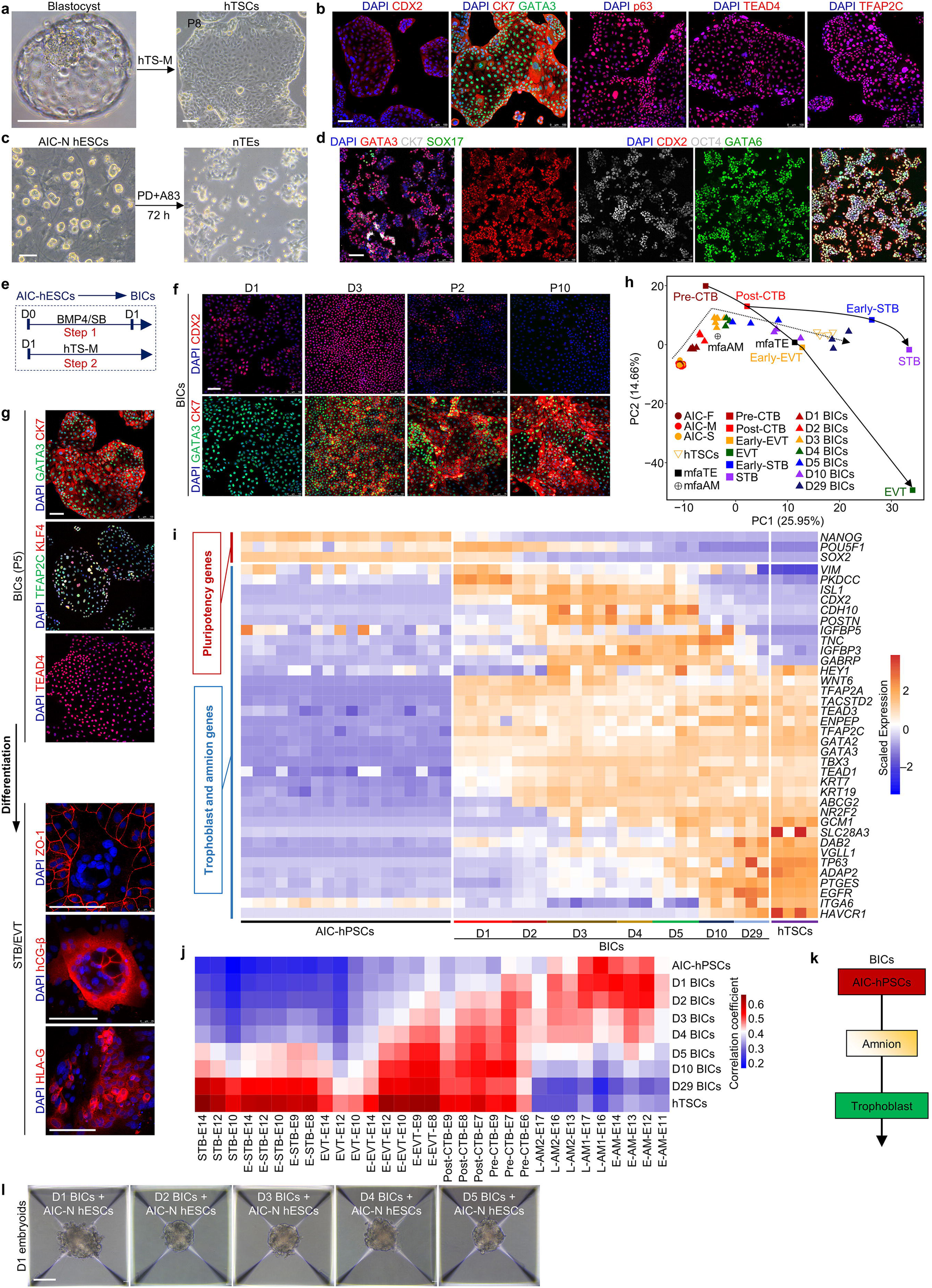
Derivation and characterization of extraembryonic cells, related to Fig. 3. **a** Representative contrast-phase images showing the derivation of human trophoblast stem cells (hTSCs) from a blastocyst ^43^. **b** Immunostaining of hTSCs with indicated markers. **c** AIC-N hESCs are induced to differentiate into nTEs (naive hESC-derived trophectoderm-like cells) in PD+A83 for 3 days ^41^. **d** Immunostaining of nTEs with indicated markers. **e** Schematic of BIC (BMP4-induced cell) generation from AIC-hESCs ^35^. SB, SB431542; hTS-M, hTSC medium. **f** Immunostaining of BICs with indicated markers at the different time points. **g** Immunostaining of BICs at passage 5 (P5) with indicated markers (top), and differentiation of BICs at P5 into syncytiotrophoblast– and extravillous trophoblast-like cells (bottom). **h** PCA analysis of bulk RNA-seq (AIC-hPSCs ^35^, hTSCs and BICs different days after induction) and single-cell RNA-seq (trophoblast cells from 3D-cultured human embryos ^4^, trophectoderm and amnion cells from monkey embryos cultured *in vitro* ^30^) data. We filtered the most variable genes among different cells of human embryo with average log_2_ (FPKM + 1) > 2 and the square of coeffificient of variation (log CV^2^) > 0.2, and 600 most variable genes expressed in both human and monkey embryos were used to perform PCA analysis. **i** Heatmap of representative pluripotency, amnion and trophoblast genes in AIC-hPSCs ^35^, BICs different days after induction and hTSCs. Values represent log_2_ (FPKM+1) scaled by gene expression across samples. AIC-hPSCs contain six cell lines (H9/hES1/S4/h1/h2/h3) cultured in the AIC medium from published data ^35^. hTSCs contain two cell lines (hTS1 and hTS2) cultured on feeders or Col IV. BICs are derived from AIC-hESCs cultured on Matrigel (H9/hES1/h1). **j** Heatmap showing Pearson correlation coefficients for expression of variable genes among human trophoblast cells ^4^, monkey amnion cells^30^, AIC-hPSCs, BICs different days after induction and hTSCs. Data of human trophoblast cells and monkey amnion cells are based on average expression level of these single cells. 2035 filtered genes were used as input for Pearson correlation analysis. STB, syncytiotrophoblast; CTB, cytotrophoblast; EVT, extravillous cytotrophoblast; AM, amnion. **k** BICs undergoes a state transition from pluripotency to amnion-like and finally to trophoblast-like fate. **l** Representative contrast-phase images of embryoids assembled from AIC-N hESCs and BICs different days after induction. A large number of dead cells surround the embryoid assembled from D1 BICs and AIC-N hESCs. Scale bars, 100 µm.

**Fig. S4.**
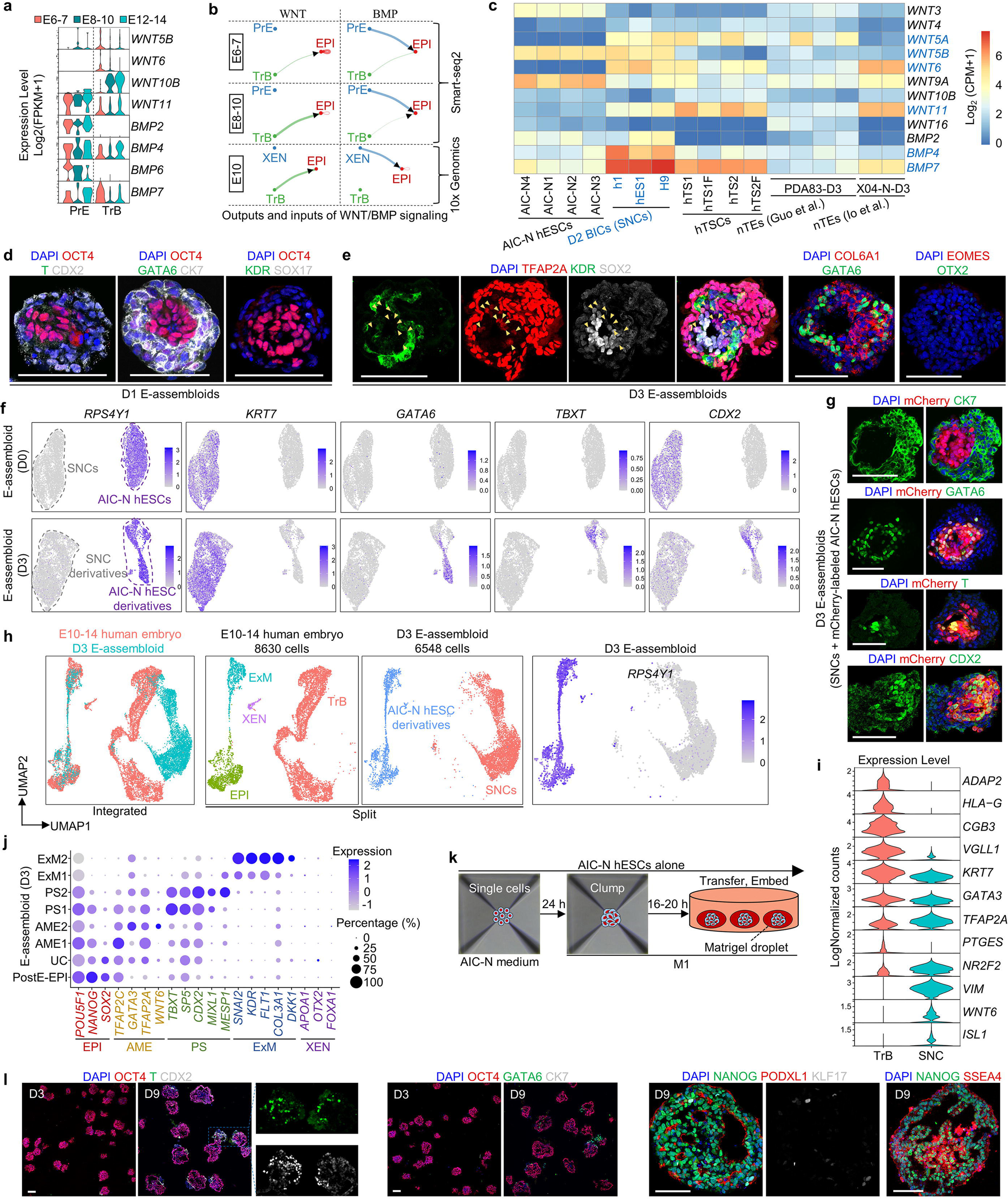
SNCs induce the development of AIC-N hESCs in E-assembloids, related to Fig. 3. **a** Violin plots of WNT and BMP ligand genes in PrE and TrB by scRNA-seq data (smart-seq2) ^4^. **b** The inferred WNT and BMP signaling outputs and inputs from extraembryonic to embryonic tissues in 3D-cultured human embryos before E10, line weight represents the communication probability. **c** Heatmap of log-transformed expression levels of major ligand genes contributing to WNT and BMP signaling among AIC-N hESCs, and D2 BICs (SNCs), hTSCs cultured on Col IV (hTS1 and hTS2) or feeders (hTS1F and hTS2F), and nTEs (PDA83-D3 ^41^ and X04-N-D3 ^42^). Values are presented as log_2_ (CPM+1). D2 BICs expressed different levels of *WNT5A/B*, *WNT6*, *WNT11*, *BMP4* and *BMP7*. D2 BICs were used to construct E-assembloids and hereinafter collectively referred to as signaling nest cells (SNCs). **d** and **e** Immunostaining of D1 and D3 E-assembloids cultured in the M1 condition with indicated markers. Yellow arrowheads indicate TFAP2A^weak^SOX2^weak/–^ AMELCs surrounded by SOX2^−^KDR^+^ ExMLCs. **f** UMAP plots showing expressions of Y-chromosome specific gene *RPS4Y1* and lineage marker genes according to scRNA-seq data from D3 E-assembloids grown in the M1 condition. The E-assembloids were constructed with male AIC-N2 hESCs and female H9 SNCs, and the scRNA-seq data from AIC-N2 hESC clumps and H9 SNCs were used as D0 control. **g** Immunostaining images of D3 E-assembloids grown in the M1 condition showed that CK7 is almost exclusively expressed in mCherry^−^ SNC derivatives, whereas GATA6, T and CDX2 are almost exclusively expressed in mCherry^+^ AIC-N hESC derivatives. E-assembloids are derived from H9 SNCs and mCherry-labeled AIC-N2 hESCs. **h** UMAP visualization and plots of integration analysis of scRNA-seq data from human embryos and D3 E-assembloids. *RPS4Y1* gene labels male AIC-N2 hESC derivatives. **i** Violin plots of indicated genes expressed in TrB (embryo) and SNC (E-assembloid) population by scRNA-seq data (10X Genomics). **j** Dot plots of candidate genes specific for the indicated cell subtypes from D3 E-assembloids. **k** and **l** Diagram (**k**) and representative staining images (**l**) that AIC-N hESC clumps were cultured alone until D3 and D9 in the M1 condition. EPI, epiblast; XEN, extraembryonic endoderm; ExM, extraembryonic mesoderm; TrB, trophoblast; SNC, signaling nest cell; PostE−EPI, post-implantation early EPI; UC, undefined cell-type; AME, amniotic epithelium; PS, primitive streak. Scale bars, 100 µm.

**Fig. S5.**
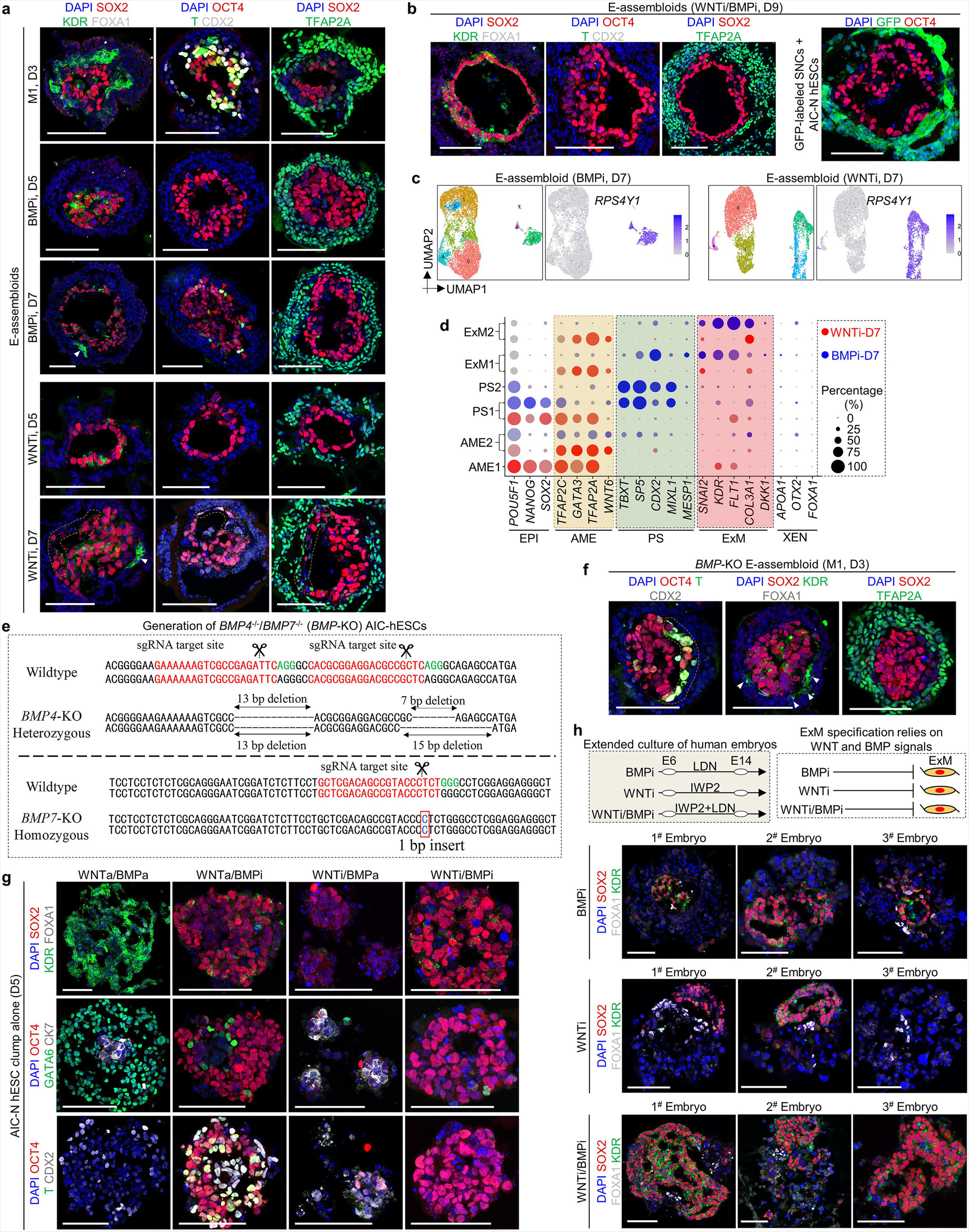
Functions of WNT and BMP signaling, related to Fig. 4. **a** and **b** Representative staining images of E-assembloids grown in indicated conditions at indicated time points. Red and white arrowheads indicate PSLCs and ExMLCs, respectively. Dotted lines show AMELCs. (B, right) E-assembloids assembled by AIC-N4 hESCs and GFP-labeled SNCs (from AIC-hES1) showed that almost all AIC-N hESC derivatives expressed the pluripotency marker OCT4 in the WNTi/BMPi condition on D9. **c** UMAP visualization of major subtype of cells and UMAP plots of *RPS4Y1* gene in E-assembloids cultured in indicated conditions on D7. **d** Dot plots of candidate genes specific for indicated cell subtypes in E-assembloids grown in indicated conditions. **e** Generation of *BMP4*^-/-^/*BMP*7^-/-^ H9 AIC-hESCs (*BMP*-KO H9 AIC-hESCs). **f** Representative staining images of D3 *BMP*-KO E-assembloids grown in the M1 condition. Red and white arrowheads indicate PSLCs and ExMLCs, respectively. Dotted lines show AMELCs. **g** Representative staining images of AIC-N hESC clumps cultured alone in indicated conditions for five days. **h** Schematic diagram of different culture conditions for human embryo (top left); functional diagram of WNT and BMP signaling on ExM specification (top right); and representative staining images of E14 human embryos grown in indicated conditions (bottom). Six blastocysts per group were cultured in the indicated conditions from E6-14, and three surviving embryos per group were used for immunostaining. E, embryonic day; EPI, epiblast; AME, amniotic epithelium; PS, primitive streak; ExM, extraembryonic mesoderm; XEN, extraembryonic endoderm; BMPi, BMP inhibition; BMPa, BMP activation; WNTi, WNT inhibition; WNTa, WNT activation. LDN, LDN193189-2HCl. Scale bars, 100 µm.

**Fig. S6.**
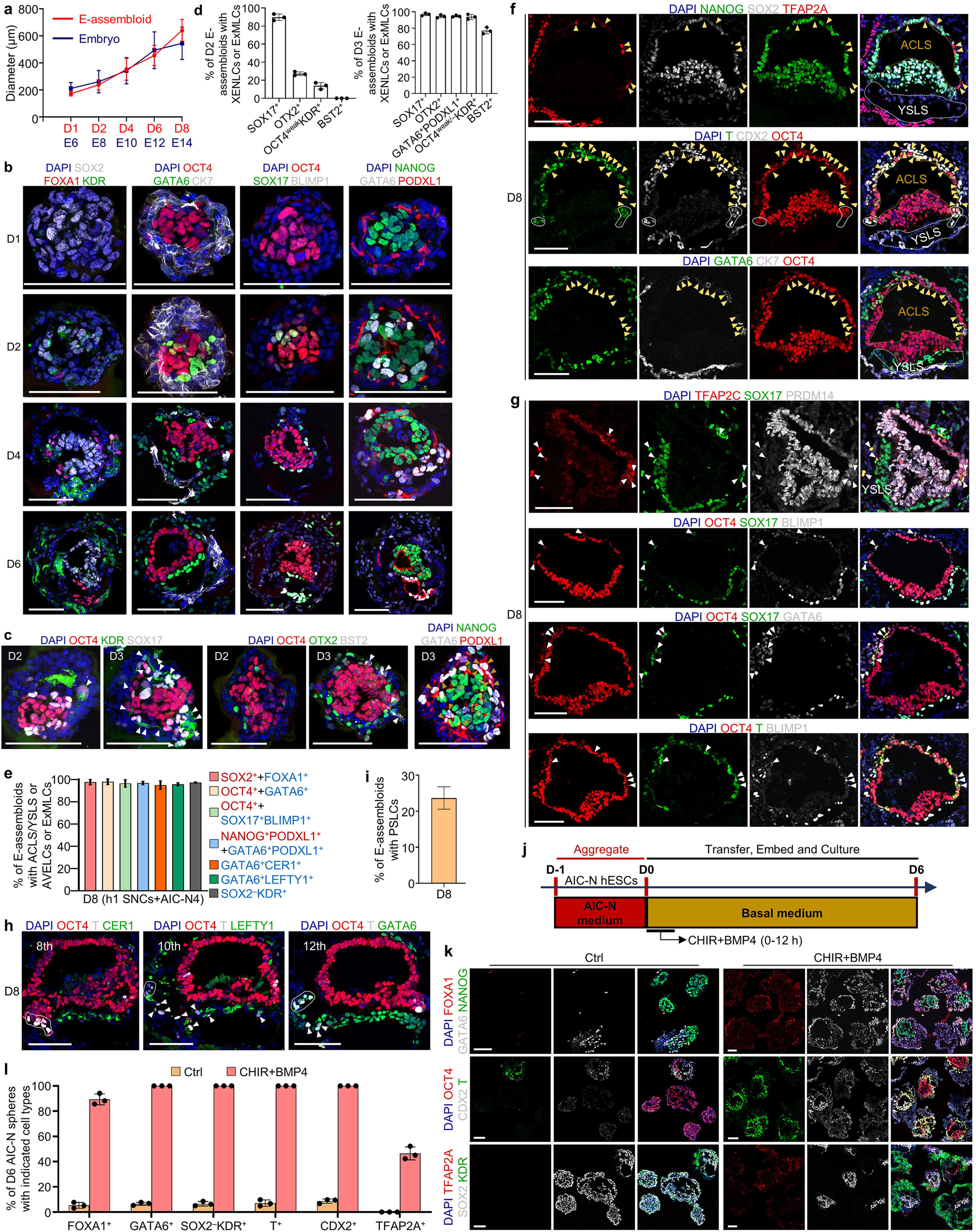
E-assembloids reconstitute 3D structures of early post-implantation embryos, related to Fig. 6. **a** Dynamic diameters of E-assembloids and 3D-cultured human embryos (published data ^4^). For E-assembloids, N=3 independent experiments; at least 100 E-assembloids were quantified each experiment; data are presented as mean±s.d. **b** and **c** Representative immunostaining images showing the dynamic development of XENLCs, ExMLCs, ACLS and YSLS in E-assembloids at different time points. **d** Quantification of the E-assembloids with XENLCs or ExMLCs on D2 and D3. Three independent experiments; more than 100 structures were quantified each experiment; data are presented as mean±s.d. **e** Repeated experiments from two other cell lines (h1 SNCs and AIC-N4 hESCs), quantification of the E-assembloids with ACLS and YSLS or AVELCs or ExMLCs on D8. N = 3 independent experiments; more than 100 structures were quantified each experiment; data are presented as mean±s.d. Red and blue represent markers of EPILCs and XENLCs/ExMLCs, respectively. **f** Representative staining images showing the specifications of AMELCs (yellow arrowheads) and PSLCs (white solid lines) in D8 E-assembloids, the yellow and white dotted lines indicate the ACLS and YSLS, respectively. **g** Representative staining images showing the specification of PGCLCs (white arrowheads) in D8 E-assembloids. **h** Representative staining images showing the generation of potential PSLCs (solid lines and arrowheads). White ordinal numbers indicate section numbers. **i** Quantification of D8 E-assembloids with putative PSLCs. N = 3 independent experiments; more than 100 structures were quantified each experiment; data are presented as mean±s.d. **j** Schematic diagram of CHIR+BMP4-treated (12 hours) AIC-N hESC clumps cultured alone for six days. **k** Representative staining images showing the generation of embryonic and extraembryonic lineages in AIC-N hESC-derived spheres. **l** Quantification of the AIC-N hESC-derived spheres with indicated cell types. Three independent experiments; more than 100 spheres were quantified each experimental group; data are presented as mean±s.d. E, embryonic day; AC, amniotic cavity; YS, yolk sac; AVE, anterior visceral endoderm; ExM, extraembryonic mesoderm; PS, primitive streak; LS, –like structure; LCs, –like cells. Scale bars, 100 µm.

**Fig. S7.**
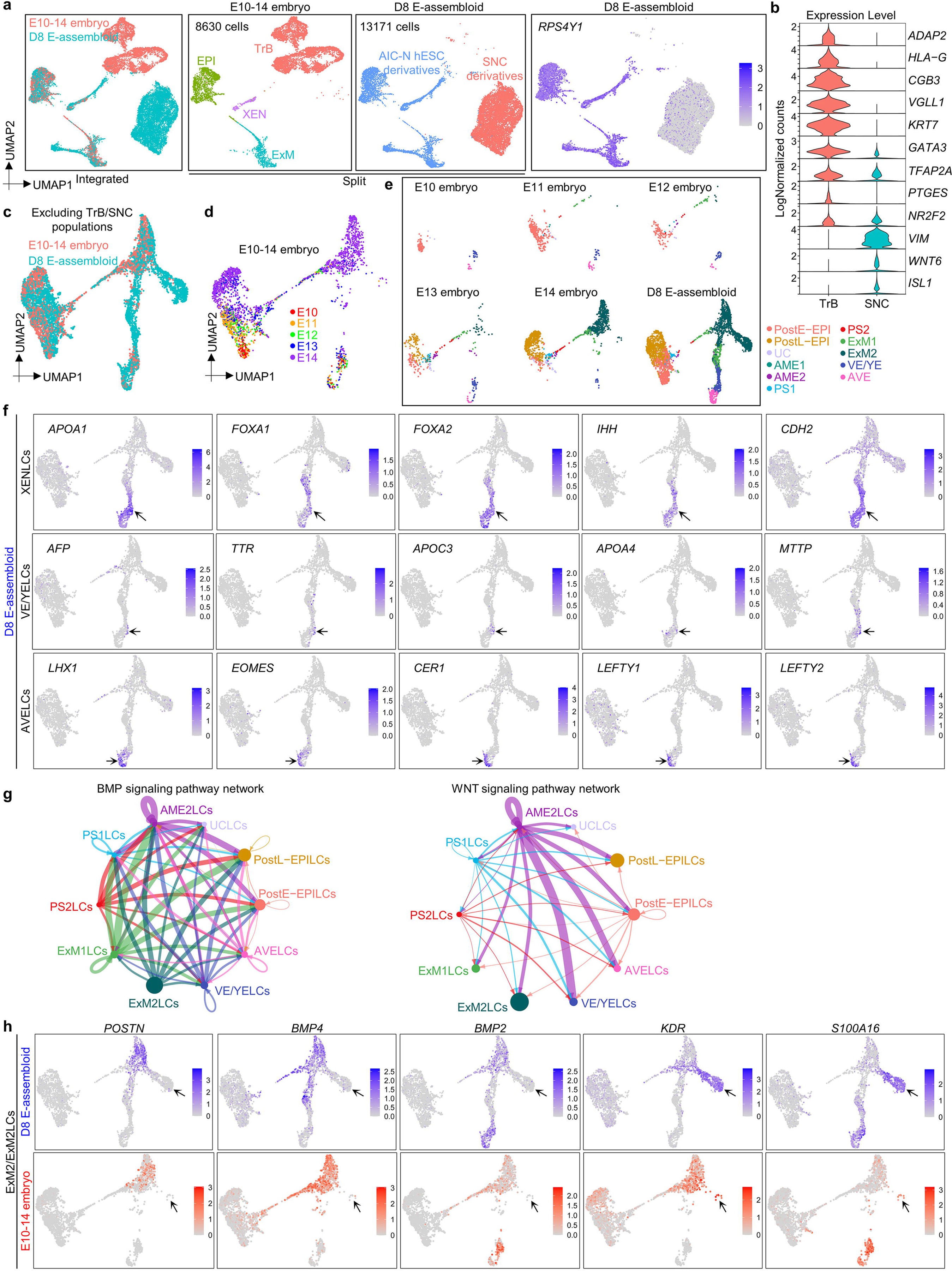
Comparing E-assembloids with 3D-cultured human embryos, related to Fig. 7. **a** UMAP visualization and plots of integration analysis of scRNA-seq data from human embryos and D8 E-assembloids. *RPS4Y1* gene labels male AIC-N2 hESC derivatives. **b** Violin plots of indicated genes expressed in TrB (embryo) and SNC (D8 E-assembloid) population by scRNA-seq data (10X Genomics). **c** UMAP visualization of integration analysis of the remaining cell types after excluding TrB and SNC population in human embryos and D8 E-assembloids, respectively. **d** UMAP visualization of the remaining cell types after excluding TrB population in human embryos at different embryonic days. **e** Split UMAP visualization of the remaining cell types after excluding TrB (E10-14 human embryos) and SNC (D8 E-assembloids) population according to different developmental time points. **f** UMAP plots of indicated genes expressed in D8 E-assembloids. From top to bottom, arrows point to XENLCs, VE/YELCs and AVELCs, respectively. **g** The inferred BMP and WNT signaling pathway networks in D8 E-assembloids. Circle sizes are proportional to the number of cells in each subpopulation and line weight represents the communication probability. **h** UMAP plots of indicated genes expressed in D8 E-assembloids and 3D-cultured embryos. Some cells in ExM2 population exhibit haemato-endothelial characteristics (arrow). EPI, epiblast; XEN, extraembryonic endoderm; ExM, extraembryonic mesoderm; TrB, trophoblast; SNC, signaling nest cell; PostE−EPI and PostL−EPI, post-implantation early and late EPI; UC, undefined cell-type; AME, amniotic epithelium; PS, primitive streak; VE/YE, visceral/yolk sac endoderm; AVE, anterior visceral endoderm; LCs, –like cells.

**Supplementary Table 1:**
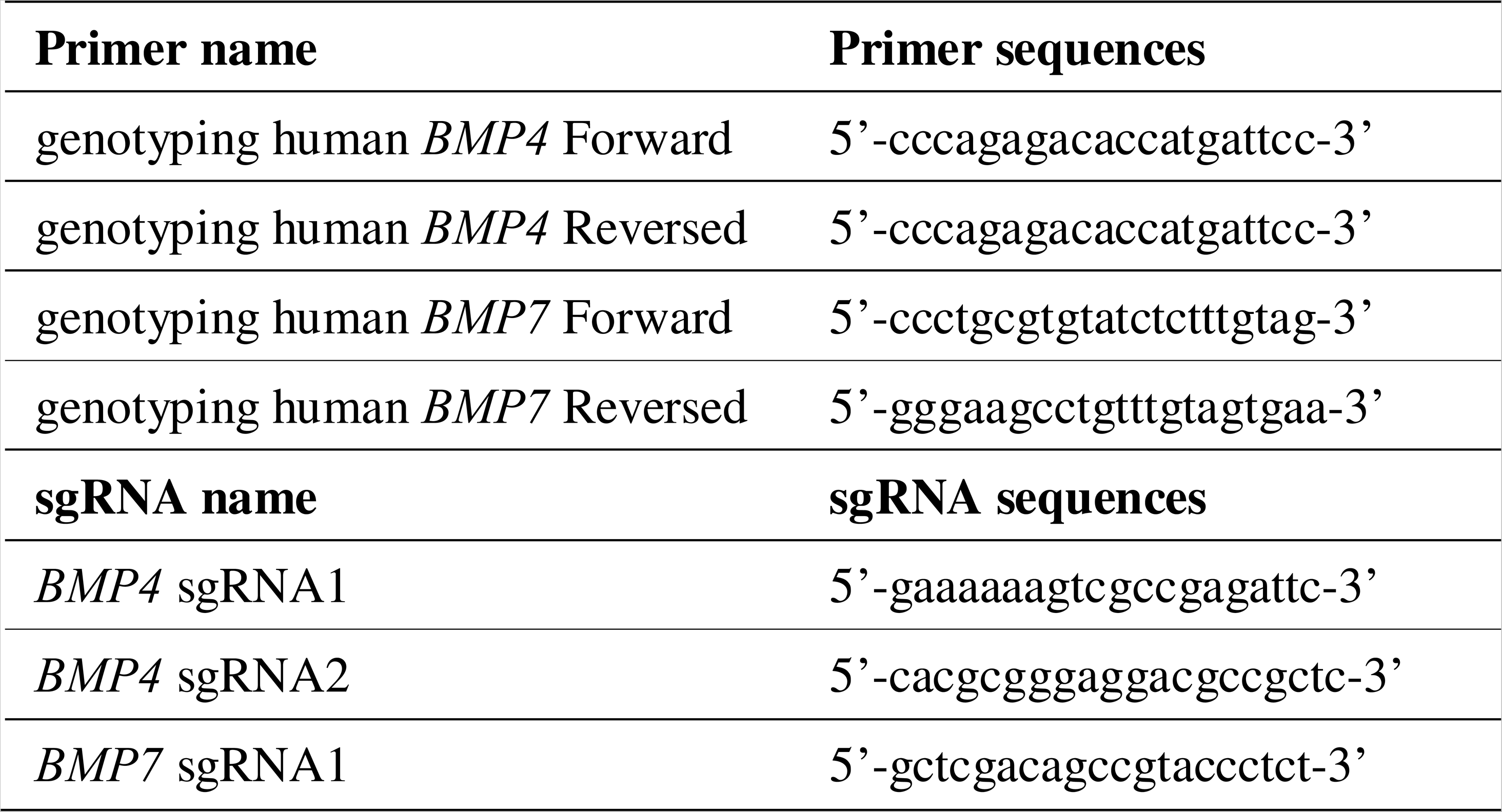
Primer and sgRNA information.

**Supplementary Table 2:** Antibody information.

| Antibody (species/isotype) | Source | Cat# | Dilution |
| --- | --- | --- | --- |
| TFCP2L1 (goat IgG) | R&D Systems | AF5726 | 1:500 |
| KLF17 (rabbit IgG) | Atlas Antibodies | HPA024629 | 1:500 |
| KLF4 (rabbit IgG) | Millipore | 09-821 | 1:600 |
| NANOG (goat IgG) | R&D Systems | AF1997 | 1:400 |
| SOX2 (rabbit IgG) | Millipore | AB5603 | 1:600 |
| OCT4 (mouse IgG) | Santa Cruz | sc-5279 | 1:600 |
| N-Cadherin (rabbit IgG) | GeneTex | GTX127345 | 1:400 |
| COL3A1 (rabbit IgG) | Abcam | Ab7778 | 1:400 |
| PRDM14 (rabbit IgG) | Millipore | AB4350 | 1:200 |
| ALPPL2 (rabbit IgG)<sup>40</sup> | Made in-house (Gao Lab) | N/A | N/A |
| β-catenin (rabbit IgG) | Cell Signaling Technology | 8480 | 1:200 |
| Tra-1-60 (mouse IgM) | Millipore | SCR001 | 1:200 |
| E-Cadherin (mouse IgG) | Abcam | ab76055 | 1:200 |
| BST2 (rabbit IgG) | Abcam | ab243230 | 1:200 |
| SUSD2-PE (mouse IgG) | Biolegend | 327406 | 1:200 |
| Tra-1-85/CD147 (rabbit IgG) | Abcam | ab108308 | 1:200 |
| GATA6 (goat IgG) | R&D Systems | AF1700 | 1:600 |
| GATA6 (rabbit IgG) | Cell Signaling Technology | 5851T | 1:600 |
| SOX17 (goat IgG) | R&D Systems | AF1924 | 1:600 |
| SOX17 (rabbit IgG) | Cell Signaling Technology | 81778 | 1:500 |
| GATA4 (goat IgG) | Santa Cruz | sc-1237 | 1:100 |
| OTX2 (goat IgG) | R&D Systems | AF1979 | 1:400 |
| FOXA2 (goat IgG) | Santa Cruz | sc-6554 | 1:200 |
| CDX2 (rabbit IgG) | Abcam | ab76541 | 1:400 |
| CK7/KRT7 (rabbit IgG) | Abcam | ab181598 | 1:500 |
| GATA3 (mouse IgG) | Thermo Fisher Scientific | MA1-028 | 1:500 |
| P63 (mouse IgG) | Abcam | ab735 | 1:100 |
| TEAD4 (rabbit IgG) | Atlas Antibodies | HPA056896 | 1:200 |
| TFAP2C (mouse IgG) | Santa Cruz | sc-12762 | 1:400 |
| HLA-G (mouse IgG) | Abcam | ab52455 | 1:200 |
| hCG-β (mouse IgG) | Abcam | ab9582 | 1:200 |
| ZO-1 (mouse IgG) | Thermo Fisher Scientific | 33-9100 | 1:100 |
| Brachyury/T (goat IgG) | R&D Systems | AF2085 | 1:600 |
| PODXL1 (mouse IgG) | R&D Systems | MAB1658 | 1:600 |
| EOMES (mouse IgG) | R&D Systems | MAB6166 | 1:200 |
| TFAP2A (mouse IgG) | Santa Cruz | sc-12726 | 1:400 |
| LAMININ (rabbit IgG) | Sigma Aldrich | L9393 | 1:100 |
| SNAIL (goat IgG ) | R&D Systems | AF-3639 | 1:400 |
| VIMENTIN (mouse IgG) | eBioscience | 14-9897 | 1:2000 |
| CER1 (rabbit IgG) | Sigma Aldrich | HPA019917 | 1:100 |
| LEFTY1 (goat IgG) | R&D Systems | AF746 | 1:500 |
| BLIMP1 (rabbit IgG) | Cell Signaling Technology | 9115S | 1:100 |
| SSEA4 (mouse IgG) | Millipore | SCR001 | 1:100 |
| CD24 (mouse IgG) | Abcam | ab134375 | 1:100 |
| FOXA1 (mouse IgG) | Abcam | ab55178 | 1:100 |
| COL6A1 (rabbit IgG) | Abcam | ab182744 | 1:400 |
| KDR (goat IgG) | R&D Systems | AF357 | 1:400 |
| Alexa Fluor 488 donkey anti-rabbit IgG | Jackson ImmunoResearch | 711-545-152 | 1:600 |
| Alexa Fluor 488 goat anti-mouse IgM | Thermo Fisher Scientific | A-21042 | 1:600 |
| Alexa Fluor 568 donkey anti-mouse IgG | Thermo Fisher Scientific | A-10037 | 1:600 |
| Alexa Fluor 488 donkey anti-mouse IgG | Thermo Fisher Scientific | A-21202 | 1:600 |
| Alexa Fluor 647 donkey anti-rabbit IgG | Thermo Fisher Scientific | A-31573 | 1:600 |
| Alexa Fluor 555 donkey anti-rabbit IgG | Thermo Fisher Scientific | A-31572 | 1:600 |
| Alexa Fluor 647 donkey anti-goat IgG | Thermo Fisher Scientific | A-21447 | 1:600 |
| Alexa Fluor 488 donkey anti-goat IgG | Thermo Fisher Scientific | A-11055 | 1:600 |

## Materials and Methods

### Ethics statement

The research about human embryo and derivation of new stem cell lines from human blastocysts is a continuation of our previous work ^4^ and had been approved by the Medicine Ethics Committee of The First People’s Hospital of Yunnan Province (2017LS[K]NO.035 and KHLL2020-KY064) ^4^. All donated embryos were supernumerary frozen embryos after IVF clinic treatment. The informed consent process for embryo donation complied with International Society for Stem Cell Research (ISSCR) Guidelines (2016/2021) and Ethical Guidelines for Human Embryonic Stem Cell research (2003) jointly issued by Ministry of Science and Technology and Ministry of Health of People’s Republic of China. All the donor couples had signed the informed consent for voluntary donations of supernumerary embryos for this study. No financial inducements were offered for the donations.

All experiments of *in vitro* culture for human E-assembloids complied with the 2016/2021 ISSCR Guidelines ^59, 60^. All E-assembloids were terminated by no later than Day 9 and did not develop beyond the appearance of primitive streak or generate the initial nervous system, meeting the internationally recognized ethical limit for human embryo model ^59, 60^. The research was approved by the Medical Ethics Committee of Yunnan Key Laboratory of Primate Biomedical Research.

### Embryo manipulation

To control the variability of the blastocysts, we evaluated the quality of the embryos before use. According to the Gardner’s scoring system ^61^, thawed blastocysts were given numerical scores from 1 to 6 based on their expansion degree and hatching status. The blastocysts with expansion and hatching status above 3 and with visible inner cell mass above grade B were used in the study. Frozen-thawed human blastocysts (5-6 days post-fertilization) were gently treated by acidic Tyrode’s solution (Sigma, T1788) to remove the zona pellucida ^4^. *In vitro* 3D culture, frozen section staining and single-cell isolation of human embryos were performed according to previously published methods ^4^. To verify the function of WNT and BMP signaling, IWP2 (2 μM, Selleck, S7085) and LDN193189-2HCl (LDN, 200 nM, Selleck, S7507) were used during 3D-culture of human embryos from embryonic day (E6-14) (Supplementary information, Fig. S5h).

In this study, 3D-cultured human embryos were selected for 10x sequencing according to previously established two criteria: 1) obvious expansion over culture; and 2) absence of obvious cell death or fragmented phenotype^4^. The numbers and the size of human embryos used for 10x sequencing are shown in Supplementary information, Fig. S1a.

### Derivation and culture of naïve hESC lines in 21% O_2_

Blastocysts were seeded on mouse embryonic fibroblast (Millipore, PMEF-CFL) feeder cells (hereafter referred to as feeders) inactivated by mitomycin C (Sigma, M0503). After 48 hours, hESC-like outgrowths became visible in the attached embryos, was mechanically transferred into 50% TrypLE (Thermo Fisher Scientific, 12605-028) and incubated for 10-15 min at 37□. Alternatively, if the hESC-like outgrowth was invisible, the whole attached embryo was incubated with 50% TrypLE for 10-15 min at 37□. Next, the digested hESC-like outgrowth (or whole embryo) was transferred into a drop of AIC-N medium and dissociated into single cells by mildly pipetting up and down with a finely drawn Pasteur pipette. All dissociated single cells were scattered onto feeders. After 3-5 days, some obvious naïve hESC colonies were observed and further dissociated into single cells by 50% TrypLE for passaging. Routinely, the AIC-N medium was changed every 2 days and the newly established AIC-N hESC lines were passaged every 3-4 days at a 1:5-1:10 split ratio. AIC-N medium was supplemented with 10 μM Y27632 (Selleck, S1049) during derivation and maintenance of AIC-N hESCs. In this study, AIC-N hESCs have been retained over 50 passages in a stable naïve state. AIC-N medium was composed of modified N2B27 medium supplemented with 10 ng/ml Activin-A (Peprotech, 120-14E), 2 μM IWP2, 0.3 μM CHIR99021 (Selleck, S2924), 1 μM PD0325901 (Selleck, S1036); 2 μM Gö6983 (TOCRIS, 2285) and 10 ng/ml recombinant human LIF (Peprotech, 300-05). 500 ml modified N2B27 medium was composed of 240 ml DMEM/F12 (Thermo Fisher Scientific, 10565-018), 240 ml Neurobasal (Thermo Fisher Scientific, 21103-049), 5 ml N2 supplement (Thermo Fisher Scientific, 17502-048), 10 ml B27 supplement (Thermo Fisher Scientific, 17504-044), 0.5% GlutaMAX (Thermo Fisher Scientific, 35050-061), 1% nonessential amino acids (NEAA, Thermo Fisher Scientific, 11140-050), 0.1 mM β-mercaptoethanol (Sigma, M7522), 0.38 ml 7.5% BSA (Sigma, A1933), 50 μg/ml L-ascorbic acid 2-phosphate (Sigma, A8960), 0.5 ml Chemically Defined Lipid Concentrate (Thermo Fisher Scientific, 11905-031), 12.5 μg/ml Insulin (Roche, 11376497001), 1.25 ml Sodium Pyruvate (Thermo Fisher Scientific, 11360-070), 0.02 μg/ml Progesterone (Sigma, P8783).

### Conversion of primed hPSCs from the conventional medium to the AIC-N medium

Conventional-cultured primed hPSCs (CC-hPSCs, KSR/bFGF medium) were digested into single cells with 50% TrypLE and inoculated at a density of 3×10^4^ cells per 3.5 cm dish onto feeders. In the AIC-N medium, CC-hESC derivatives maintained stable primed state after extended single-cell passaging (every 3-4 days at a 1:5 to 1:10 split ratio), with no obvious cell differentiation, death and pluripotent state transition. We recommend supplementing AIC-N medium with 5 μM Y27632 in the first 48h of conversion.

To reset human CC-hPSCs, single CC-hPSCs were inoculated at a density of 8×10^4^ cells per 3.5 cm dish onto feeders in AIC-N medium supplemented with 1mM Valproic acid (VPA, Sigma, P4543) ^36^. After 4-6 days, cell cultures were digested into single cells using 50% TrypLE and were re-plated onto feeders at a split ratio of 1:10 followed by withdrawal of VPA, and majority of colonies displayed domed morphology at passage 1. Compared with the flat colonies, the dome-shaped colonies were first detached from feeders by exposure to 1 mg/ml Collagenase type IV (Thermo Fisher Scientific, 17104-019). Based on the different detaching speed of colonies, we purified domed colonies in the context of Collagenase type IV during initial (1-2) passages, the detached colonies were collected and dissociated into single cells with 50% TrypLE for sub-culturing. Thereafter, cell cultures can be split every 3-4 days at a 1:5 ratio via single cell dissociation with 50% TrypLE, medium was routinely changed every other day. AIC-N medium was supplemented with 10 μM Y27632 during reset and maintenance of naïve hESCs.

### Establishment and culture of hTSC lines

Establishment and culture of blastocyst-derived hTSC lines was performed according to the published method ^43^. Briefly, blastocyst was placed into Col IV (Corning, 354233) –coated 96-well plate and cultured in 150 µL of hTSC medium. After 4-5 days of culture, the blastocyst outgrowth was dissociated with 50% TrypLE for 10-20 min at 37□ and passaged to a new Col IV-coated 96-well plate. hTSCs were routinely passaged every 3-4 days at a 1:3 split ratio and pure hTSC lines were established within 5-10 passages. hTSC medium was composed of the following ingredients: DMEM/F12 supplemented with 0.1 mM 2-mercaptoethanol, 0.2% FBS (BI, 04-002-1A), 0.3% BSA, 1% ITS-X supplement (Thermo Fisher Scientific, 51500-056), 50 μg/ml L-ascorbic acid 2-phosphate, 50 ng/ml EGF (R&D Systems, 236-EG-01M), 2 μM CHIR99021, 0.5 μM A83-01, 1 μM SB431542, 0.8 mM VPA and 5 μM Y27632. Alternatively, hTSCs can also be cultured on Feeders instead of Col IV in hTSC medium.

### Differentiation of AIC-N hESCs into nTEs and hTSCs

The differentiation of nTEs (Naïve hESC-derived trophectoderm-like cells) was performed according to previously published methods ^41, 42^, with some minor modifications. Briefly, single AIC-N hPSCs were inoculated at a density of 1×10^5^ cells per 3.5 cm dish onto Matrigel (Corning, 354277)-coated dishes in the nTE induction medium, which was changed every two days. After 72 hours, nTEs were dissociated and collected to assemble embryoids. nTE induction medium was composed of modified N2B27 medium supplemented with 2 µM PD0325901, 2 µM A83-01 and 5 µM Y27632.

The differentiation of hTSCs was performed as described ^41, 42^, with some minor modifications. Briefly, single AIC-N hPSCs were inoculated at a density of 5×10^4^ cells per 3.5 cm dish onto feeders, after 3 days treatment with 1 µM PD0325901 and 1 µM A83-01 in the modified N2B27 medium ^35^, the initial medium was replaced with hTSC medium ^43^ and the cultures were passaged at day 5. Henceforth, cells were routinely passaged every 3-4 days at a 1:3 split ratio and pure hTSC lines were established within 2-3 passages. All culture medium was supplemented with 5 μM Y27632 and replaced every two days. 500 ml modified N2B27 medium was composed of 240 ml DMEM/F12, 240 ml Neurobasal, 5 ml N2 supplement, 10 ml B27 supplement, 0.5% GlutaMAX, 1% NEAA, 0.1 mM β-mercaptoethanol, 0.38 ml 7.5% BSA, 50 μg/ml L-ascorbic acid 2-phosphate, 0.5 ml Chemically Defined Lipid Concentrate, 12.5 μg/ml Insulin, 1.25 ml Sodium Pyruvate and 0.02 μg/ml Progesterone.

### Differentiation of AIC-N hESC-derived hTSCs into STBs and EVTs

The differentiation methods of AIC-N hESC-derived hTSCs into STBs (syncytiotrophoblast cells; 2D and 3D) and EVTs (extravillous cytotrophoblast cells) were performed as previously reported ^43^.

### AIC-hPSC culture, BIC induction and terminal differentiation

AIC-hPSCs grown on Matrigel-coated dishes/plates were cultured in the AIC medium as described previously ^35^. Briefly, the AIC medium was changed every two days and AIC-hPSCs were passaged every 3-4 days at a 1:10-1:20 split ratio by single-cell dissociation. AIC medium was composed of modified N2B27 medium supplemented with 10 ng/ml Activin-A, 2 μM IWP2 and 0.6 μM CHIR99021.

For generation of BMP4-induced cells (BICs), AIC-hPSCs were digested into single cells with 50% TrypLE and inoculated at a density of 1×10^5^ cells per 3.5 cm dish onto Matrigel-coated dishes in the BIC induction medium. After 24 h, BIC induction medium was replaced with hTSC medium ^43^ and BICs were passaged at Day 4 after induction. Routinely, hTSC medium was changed every 2 days, and BICs were passaged every 3-4 days at 1:5-1:10 and expanded over 10 passages. D2 BICs were termed signaling nest cells (SNCs) in the following context. Medium was supplemented with 5 μM Y27632 during induction and maintenance of BICs. BIC induction medium is composed of DMEM/F12, 15% knockout serum replacement (KSR, Thermo Fisher Scientific, A3181502), 1% NEAA, 0.1 mM β-mercaptoethanol and 12.5 μg/ml Insulin supplemented with 10 ng/ml BMP4 and 8 μM SB431542.

The differentiation of BICs into mature trophoblast cells was performed according to the published method ^62^. Briefly, BICs were passaged using 50% TrypLE and replated on Matrigel-coated plates in feeder-conditioned medium (DMEM/F12 containing 15% KSR, 1% NEAA, 0.1 mM β-mercaptoethanol and 12.5 μg/ml Insulin was pre-conditioned on feeders for 24 h, termed FCM) supplemented with 10 ng/mL BMP4, 8 μM SB431542 and 5 μM Y27632. Routinely, the medium (termed FCM+BMP4/SB) was changed every two days. These cells were fixed and stained at Day 6.

### Specifications of extraembryonic endoderm (XEN) and extraembryonic mesoderm (ExM)

For the functional experiments of WNT, BMP and Nodal signals on specifications of XENLCs and ExMLCs, AIC-N hESCs were plated at 1500/well-96 onto feeders in indicated differentiation conditions (Fig. 5c, f). The medium was changed every other day and the cells were fixed and stained at Day 4. The differentiation medium was composed of modified N2B27 medium supplemented with 50 ng/ml recombinant human FGF4 (Peprotech, 100-31-25), 1 μg/ml heparin (Sigma, H3149) and 10 μΜ Y27632. 2 μΜ CHIR99021, 10 ng/ml BMP4 (R&D Systems, 314-BP-050), 0.5 μM A83-01 (Tocris, 2939), 1 μM SB431542 (Cellagen, C7243), 2 μM IWP2 and 200 nM LDN were added to the differentiation medium according to indicated combinations in Fig. 5c, f.

### Derivation of AAVS1-Knockin hESC lines

2.5×10^5^ hESCs (AIC-hESCs or AIC-N hESCs) were electroporated with 1 μg of donor plasmid AAVS1-CAG-hrGFP (Addgene, #52344) or AAVS1-Pur-CAG-mCherry (Addgene, #80946), and 1 μg of sgRNA-CAS9 expression vector by 4D-Nucleofector (Lonza) using P3 Primary Cell 4D-Nucleofector X kit (Lonza). Transfected cells were plated onto irradiated DR4 MEFs (the Cell Bank of the Chinese Academy of Sciences, Shanghai, China; https://www.cellbank.org.cn) in AIC or AIC-N media. Three days later, cells were selected with puromycin (0.25 μg/ml) and expanded in the same media for 2 weeks, GFP/mCherry-positive and puromycin-resistant clones were picked and identified by PCR.

### Generation of AIC-hESCs with simultaneous knockout of *BMP4* and *BMP7*

Pre-validated gRNA sequences targeting the exon 3 of *BMP4* or *BMP7* gene were obtained from genome-wide databases provided by GenScript (https://www.genscript.com/gRNA-database.html) ^63^, and cloned into the pGL3-U6-2sgRNA-ccdB-EF1a-Cas9ZF-IRES (Addgene, Plasmid#115519) vector. 2 μg of assembled pGL3 plasmid were electroporated into 2.5×10^5^ AIC-H9-hESCs ^35^ by 4D-Nucleofector (Lonza) using P3 Primary Cell 4D-Nucleofector X kit (Lonza). Transfected cells were plated onto irradiated DR4 MEFs (the Cell Bank of the Chinese Academy of Sciences, Shanghai, China; https://www.cellbank.org.cn) in the AIC medium ^35^. After 24 hours, cells were selected with puromycin (0.25 μg/ml) for 3 days. About one week later, the single-cell derived clones were picked for genotype analysis around the sgRNA targeting site by genomic PCR and Sanger sequencing. The PCR Primer information and the sgRNA information are in Supplementary Table 1.

### Teratoma formation

Teratoma assay was performed according to NIH guidelines and animal procedures and approved in advance by the Animal Care and Use Committee of Yunnan Key Laboratory of Primate Biomedical Research. Approximately 10^6^ cells were resuspended in 75 μl of AIC-N medium including 20 μM Y27632 and co-injected subcutaneously with 75 μl of Matrigel (Corning, 354234) into the groin of 4-week-old NOD/SCID female SPF mice (*Prkdc^scid^*/NcrCrl, Beijing Vital River Laboratory Animal Technology Co., Ltd, http://www.vitalriver.com/proinfo.aspx?nid=54). After 10-12 weeks, teratomas were excised, fixed with 4% PFA, sectioned and stained with hematoxylin and eosin.

### G-banding Karyotype Analysis

Cells were used to perform karyotype analysis one day before passaging. After incubated for 2-4 hours with fresh medium, cells were treated by Colcemid Solution (BI, 12-004-1D) at a final concentration of 0.02 μg/ml for 1 h. The cells were washed twice in PBS, trypsinized into single cells and centrifuged. Next, the pellet was resuspended in 5 ml of hypotonic solution (0.075 M KC1) and left at 37□ for 15 min. 1 mL of ice-cold fixative (3:1 methanol: acetic acid) was added dropwise to the hypotonic suspension and left at room temperature for 5 min. After spinning and removing the supernatant, 5 ml of ice-cold fixative was added dropwise to the suspension, left at room temperature for 30 min and spun down. The fixing step was repeated for three times. Finally, the pellet was resuspended in a final volume of 3 ml ice-cold fixative and placed in –40□ freezer. Subsequent G-banding karyotype analysis was performed at The First People Hospital of Yunnan Province, Kunming, China. For each analysis, at least 20 metaphases were counted.

### Blastoid formation

Blastoid formation was performed according to a previously described protocol^17^ with minor modifications. Briefly, AIC-N hESC colonies were detached from feeders by exposure to 1 mg/ml Collagenase type IV for 60-90 min. The detached colonies were collected, dissociated into single cells with 50% TrypLE, filtered through a 20 μm cell strainer (Miltenyi Biotec, 130-101-812), pelleted by centrifugation for 4 min at 1000 rpm, suspended in modified N2B27 medium containing 10 µM Y-27632 and counted using a hemocytometer. AggreWell™ 400 24-well Plates (STEMCELL Technologies, 34415) were prepared following the manufacturer’s instructions. 2 ml of cell suspension (including 8×10^4^ AIC-N hESCs) per well was added into the Aggrewells. After 24 hours, the modified N2B27 medium containing 10 µM Y-27632 was replaced with modified PALLY medium ^17^. The modified PALLY medium was composed of modified N2B27 medium supplemented with 1 µM PD0325901, 1 µM A83-01, 4 µM 1-Oleoyl lysophosphatidic acid sodium (LPA; Sigma, L7260), 10 ng/ml recombinant human LIF and 10 μΜ Y27632. After 72 hours, the modified PALLY medium was replaced with modified N2B27 medium containing 4 µM LPA and 10 µM Y-27632 and maintained for another two days. At day 5, blastoids were collected for staining. Cultures were maintained in a humidified incubator under 37°C, 5% CO_2_ and 5% O_2_ along the whole process, and the medium was refreshed every 24 hours.

### Generation of human embryoids

AIC-N hESC colonies were detached from feeders by exposure to 1 mg/ml Collagenase type IV for 60-90 min. The detached colonies were collected, dissociated into single cells with 50% TrypLE, filtered through a 20 μm cell strainer, pelleted by centrifugation for 4 min at 1000 rpm, suspended in AIC-N medium and counted using a hemocytometer. AggreWell™ 400 24-well Plates were prepared following the manufacturer’s instructions. 500 μl of cell suspension (including 3×10^4^ AIC-N hESCs) per well was added into the Aggrewells. After 24 h, AIC-N hESCs formed tight and round clumps. Different types of extraembryonic cells (xEMs, including hTSCs, nTEs and BICs/SNCs) were dissociated into single cells, filtered, centrifuged, suspended in M1, and counted as described above. AIC-N medium were largely removed and 1 ml of cell suspension (including 3.5×10^5^ xEMs) per well was slowly added to the Aggrewells. The time of cell seeding is considered as the starting point (0 h). After 1 h, additional 1 ml M1 with 2% Matrigel (Corning, 354230) per well was slowly added into the Aggrewells. Within 16-20 h, xEMs aggregated with AIC-N hESC clumps to form aggregates (embryoids). The embryoids assembled by AIC-N hESC clumps and SNCs (D2 BICs) were hereinafter collectively referred to as embryo-like assembloids (E-assembloids). The ingredients of M1 medium were as follows: 3:1 mixture of DMEM/F12 and Neurobasal supplemented with 0.1 mM 2-mercaptoethanol, 0.5% NEAA, 0.1% FBS, 2.5% KSR, 0.25% N2 supplement, 0.5% B27 supplement, 0.15% BSA, 0.25% GlutaMAX, 1% ITS-X supplement, 50 μg/ml L-ascorbic acid 2-phosphate, 50 nM Methylene blue (Selleck, S4535), 25 ng/ml EGF, 1 μM CHIR99021, 0.25 μM A83-01, 0.5 μM SB431542, 0.4 mM VPA. Additional chemicals described in this study contain: IWP2 (2 μM) and LDN (200 nM). All media were supplemented with 10 μM Y27632 during generation and culture of human embryoids.

### ‘3D embedded’ extended culture of human E-assembloids

Human E-assembloids were transferred from the microwells into the precooled 1.5-mL micro-centrifuge tubes inserted in ice. After about 10 min, the supernatant was aspirated, and ice-cold 1:2 mixture of M1 and Matrigel (Corning, 354230) was added to resuspend the pellet at a final concentration of 600 E-assembloids per ml. After mixed thoroughly (To ensure the uniform distribution of E-assembloids in the suspension, each 1.5-mL micro-centrifuge tube contained no more than 300 μL of suspension, which was thoroughly mixed while being dropped to 3.5cm dish), 10-20 μl/droplet E-assembloid suspension was plated into 3.5 cm dish (about 20 droplets per dish), the dish was quickly turned upside down to prevent E-assembloids from falling to the bottom of the droplet, allowed to solidify at 37□ for 20 min and overlaid with 2 ml pre-warmed M1 per dish. Matrigel, media and tubes were kept on ice during the whole process of embedding, and all media used for embedding and culturing human E-assembloids were supplemented with 10 μM Y27632. Routinely, the medium was changed every two days unless otherwise noted. For the extended culture of human E-assembloids, IWP2/LDN addition and CHIR/A83-01/SB431542 withdrawal were performed based on the M1 condition during indicated timeframe, and two protocols for assembly and extended culture of E-assembloids were established (Fig. 3a, d and Fig. 6a). For the functional experiments of WNT and BMP signaling, IWP2/LDN addition and CHIR withdrawal were performed based on the M1 condition (Fig. 4a).

### ‘3D embedded’ culture of AIC-N hESC clumps

AIC-N hESC clumps was prepared as described in ‘Generation of human embryoids’. After the formation of clumps, the AIC-N medium was largely removed and 2 ml of indicated medium per well was gently added to the Aggrewells. Within 16-20 h, these AIC-N hESC clumps were transferred from the microwells and cultured according to the ‘3D embedded’ culture method as described above. Briefly, these AIC-N hESC clumps were transferred into the precooled 1.5-mL micro-centrifuge tubes inserted in ice. After about 10 min, the supernatant was aspirated, and ice-cold 1:2 mixture of indicated medium and Matrigel was added to resuspend the pellet with a final concentration of 1000 clumps/ml. 10-20 μl/droplet AIC-N hESC clump suspension was plated into 3.5 cm dish (about 20 droplets per dish), allowed to solidify at 37□ for 20 min and overlaid with 2 ml pre-warmed medium per dish. Matrigel, media and tubes were kept on ice during the whole process of embedding, and all media used for embedding and culturing AIC-N hESC clumps was supplemented with 10 μM Y27632. Routinely, the medium was changed every two days unless otherwise noted. During extended culture, AIC-N hESC clumps were cultured alone to different time points in M1 condition (Supplementary information, Fig. S4k). For the functional experiments of WNT and BMP signaling, CHIR99021 (3 μM), BMP4 (20 ng/ml), LDN (200 nM) and IWP2 (2 μM) were added to M1 medium minus CHIR99021 according to indicated combinations (Fig. 4g).

### Inducing ‘3D embedded’ AIC-N hESC clumps using CHIR99021 and BMP4

AIC-N hESC clumps was prepared as described in ‘Generation of human embryoids’. 24 h after seeding, these formed AIC-N hESC clumps were transferred from the microwells and cultured for 6 days in the indicated condition (Supplementary information, Fig. S6j) according to the ‘3D embedded’ culture method as described above. The basal medium was composed of modified N2B27 medium supplemented with 10 μΜ Y27632. For the induction of WNT and BMP signaling, CHIR99021 (2 μΜ) and BMP4 (1 ng/ml) were used to treat the AIC-N hESC clumps during the first 12 h of extended culture (Supplementary information, Fig. S6j).

In this study, we usually used AIC-N hESCs at passages 20-40 to construct embryoids and did not observe obvious differences caused by the number of passages. Unless otherwise specified, cell or embryoid culture experiments were performed in a humidified incubator under 21% O_2_ and 5% CO_2_ at 37□. Cell lines were routinely checked for mycoplasma contaminations using MycoAlert Mycoplasma Detection Kit (LONZA, LT07-318) every two weeks, and all cell samples used in this study have been ruled out of mycoplasma contamination.

### Flow Cytometry

To detect cell-surface markers using flow cytometry, AIC-N hESC colonies were detached from feeders by exposure to 1 mg/ml Collagenase type IV for 60-90 min, the detached colonies were collected and dissociated into single cells with 50% TrypLE, centrifuged, washed with ice-cold PBS containing 1% FBS (BI, 04-001-1A). For flow cytometric analysis of Tra-1-85, the cells were incubated at 4□ for 30 min with an anti-Tra-1-85 antibody (Abcam, ab108308, 1:100) diluted in PBS including 1% BSA and then stained with a AlexaFluor 555-conjugated donkey anti-rabbit IgG secondary antibody (Thermo Fisher Scientific, A-31572, 1:1000) for 30 min. For Tra-1-60 and SUSD2, the cells were incubated at 4□ for 30 min with a PE-conjugated anti-Tra-1-60 antibody (Thermo Fisher Scientific, 12-8863-82, 1:5) or a PE-conjugated anti-SUSD2 antibody (Biolegend, 327406, 1:100) diluted in PBS including 1% BSA. Normal rabbit IgG (Abcam, ab172730, 1:100), PE-conjugated mouse IgM (Thermo Fisher Scientific, MGM04, 1:10), and PE-conjugated mouse IgG1 (BD Pharmingen, 554680, 1:100) antibody was used as isotype control, respectively. Flow cytometry was performed using a FACSAria III, and the data were analyzed using FlowJo software.

### Immunofluorescence staining

All adherently growing cells in the study were fixed with 4% paraformaldehyde (PFA) for 20 min at room temperature and washed thrice in PBS. For embryoids at D1 (aggregates in Aggrewells), aggregates were transferred from the microwells into the precooled 1.5-mL micro-centrifuge tubes inserted in ice. After about 10 min, the supernatant was aspirated, aggregates were resuspended and fixed in 4% PFA at 4□°C for 3 h. For ‘3D embedded’ culture, embryoids or hESC spheres were fixed in 4% PFA at 4□°C for 3 h, then transferred into the 1.5-mL micro-centrifuge tubes. All structures (embryoids or hESC spheres) were washed thrice in PBS, dehydrated overnight in PBS including 20% sucrose at 4□, embedded into O.C.T. (Sakura Finetek, 4583) and sectioned by a Leica frozen slicer at a thickness of 10 μm. After permeabilization and blocking with PBS including 0.2% Triton X-100, 100 mM Glycin and 3% BSA at room temperature for 60 min, the cells or sections were incubated with primary antibodies at 4□°C overnight, washed thrice with PBS including 0.05% Tween-20, incubated with secondary antibodies for 2 h at room temperature and washed thrice with PBS including 0.05% Tween-20. DAPI (Roche Life Science, 10236276001) was used for staining the nuclei. Pictures were taken by Leica SP8 laser confocal microscope. The antibodies were listed in Supplementary Table 2.

### Envelopment efficiency evaluation, quantifications of different types of cells or structures, cell number counts and diameter measurements

To evaluate the efficiency of AIC-N hESC clumps enveloped by xEMs (hTSCs or nTEs or BICs/SNCs), frozen sections of embryoids were stained with OCT4 and CK7 for AIC-N hESCs and xEMs, respectively. According to the expression pattern of lineage markers, these structures are divided into three types (Fig. 3b). The percentage of different types of structures were quantified manually using the confocal microscope (Leica SP8). Statistical analysis and plotting were performed with GraphPad Prism 9.

To quantify the proportion of different types of cells or embryoids, and number of specific cell types in embryoids, 2D cell cultures and frozen sections of different types of embryoids were stained with indicated markers. The percentage of different types of cells or embryoids, and number of specific cell types in embryoids were then quantified manually using the confocal microscope (Leica SP8). Statistical analysis and plotting were performed with GraphPad Prism 9.

To measure the diameters of developing E-assembloids, E-assembloids at different time points were fixed in 4% PFA, and randomly photographed by phase contrast microscope after Matrigel was depolymerized. Images were processed and diameters of E-assembloids were measured with Image J software. Statistical analysis and plotting were performed with GraphPad Prism 9.

### RNA sequencing and data analysis

Adherently growing cells (BICs grown on Matrigel and hTSCs grown on Col IV) were collected by dissociating into single cells with 50% TrypLE. For AIC-N hESCs and hTSCs grown on feeders, the colonies were detached from feeders by exposure to Collagenase type IV for 60-90 min, and the detached colonies were collected. For differentiated AIC-N hESC clumps embedded in Matrigel droplets, the cultures were subjected to Cell Recovery Solution (Corning, 354253) at 4□ for 1 h. After Matrigel was depolymerized, the differentiated AIC-N hESC clumps were collected. All cultures were washed twice in PBS. Total RNA of AIC-N hESCs, hTSCs, BICs and differentiated AIC-N hESC clumps was isolated with the TRIzol™ Reagent (Thermo Fisher Scientific, 15596018). The 2×150 bp paired end libraries were sequenced with Illumina HiSeq X Ten instrument. Library construction and sequencing were performed by Annoroad Gene Technology (http://www.annoroad.com/). The published data used in this study are from reset hPSCs ^33^, 3iL hPSCs ^64^, human EPS cells ^65, 66^, 5i/LAF hPSCs ^67^, AIC-hPSCs ^35^, NHSM4i hPSCs ^68, 69^, HENSM/HENSM-ACT hPSCs ^34^, HNES cells ^70^, human rsESCs ^71^ and conventional (primed) hPSCs. Reads were mapped to human genome hg38 using HISAT2 (v2.2.1) with GENECODE v30. The counts and FPKM values for each gene were calculated with tringTie (v2.1.1). We identified the most variable genes through fitting a non-linear regression curve between average log_2_ (FPKM) and the square of coefficient of variation according to the methods described ^41, 54^. Principal components analysis was performed using princomp function from the R stats package based on the covariance matrix. Heatmaps were generated using pheatmap package from the R software.

Correlation analysis among human trophoblast cells ^4^, monkey amnion cells ^30^, AIC-hPSCs ^35^, BICs different days after induction and hTSCs were performed by Pearson correlation. We calculated the average gene expression level of different cell types using the AverageExpression function in the Seurat. Then, specifific thresholds were applied along the x-axis (average log_2_ FPKM > 5) and y-axis (log CV^2^ > 0.2) to identify the most variable genes. Finally, the remaining genes are used to calculate the Pearson correlation among different cell types.

### Bisulfite sequencing and data analysis

Paired-end sequencing was performed on HiSeq X Ten platform (Illumina). Library construction and sequencing were completed by Annoroad Gene Technology (http://www.annoroad.com/).

The sequencing quality of paired end 150bp raw reads were checked by FastQC (v0.11.8) (https://www.bioinformatics.babraham.ac.uk/projects/fastqc/). Reads were then aligned to human reference genome (GRCh38) using BSMAP (v2.7.4) allowing 2 mismatches (-v 2), up to one multiple hits (-w 1), map to all four possible strands (-n 1) ^72^. Methylation levels over each cytosine were calculated using BSMAP (v2.7.4) methratio.py script. Methylation levels over different genomic regions were extracted using a customized Python script (https://github.com/MoPie/20210607LTQProject/extractMeth.py). Promoters were defined as upstream 1kb and downstream 1kb of transcription start sites. CpG islands (CGIs) coordinates were downloaded from UCSC genome browser (http://hgdownload.cse.ucsc.edu/goldenpath/hg38/database/cpgIslandExt.txt.gz). Promoter CGIs on chromosome X were calculated using Bedtools intersect function with default parameter ^73^. Promoter non-CpG islands were defined as promoter with no CGIs. Random 2kb bins were permutated with Bedtools shuffle function by excluding CGIs and promoter regions ^73^. Data were then visualized in R with boxplot() function and pheatmap() function from pheatmap package (https://cran.r-project.org/web/packages/pheatmap/). Global comparison of methylation between samples was calculated by averaging CpG methylation levels over indicated windows using the bisulfite feature methylation pipeline in SeqMonk. Pseudocolour scatter plots of methylation levels over indicated windows were generated using R.

### Single cell dissociation, RNA sequencing and data processing

Single cell dissociation of 3D-cultured human embryos was optimized based on our previous work and other report ^4, 74^. Briefly, embryos were washed in PBS three times, washed with TrypLE two times and incubated in TrypLE at 37□ for 25-30 min. The embryos were dissociated into single cells by mouth pipetting in 1% DFBS/PBS. E-assembloids grown in indicated conditions at different time points were subjected to Cell Recovery Solution at 4□ for 1 h. After Matrigel was depolymerized, E-assembloids were transferred into TrypLE containing 1 mg/ml Dispase (Corning, 354235) and incubated for 30 min at 37°C. E-assembloids were gently dissociated into single cells by pipetting up and down, filtered through a 20 μm cell strainer, centrifuged, suspended in PBS containing 0.04% BSA and counted using a hemocytometer. Single cell suspension of E-assembloids and embryos were loaded into the 10x Genomics Chromium system within 30 minutes after dissociation. 10x Genomics v3 or v3.1 libraries were prepared according to the manufacturer’s instructions. Libraries were sequenced with a minimum coverage of 30 000 raw reads per cell on an Illumina HiSeq X Ten with 150-bp paired-end sequencing, which was performed by Annoroad Gene Technology (http://www.annoroad.com/). Sequencing data was aligned and quantified using the Cell Ranger Pipeline v3.0.1 (10x Genomics) against the GRCh38 reference genome. Data from 10x Genomics for E-assembloids or 3D-cultured embryos were filtered based on number of expressed genes and expression level of mitochondrial genes (below 20%). Cell doublets were removed by DoubletFinder (v2.0.3) with assuming multiplet rate according to the loaded cells number (refer to Multiplet Rate Table provided in the 10x Genomics User Guide).

Further analyses were performed using Seurat package (v 4.0.3) ^75^. The raw counts were normalized and scaled with default parameters. Top 2000 most variable genes were identified and used for dimensionality reduction with PCA and followed with non-linear dimensionality reduction using UMAP. Cell types were defined based on the lineage markers and clusters identified through FindClusters function. Data was visualized with the UMAP dimensionality reduction. DEGs were identified with the FindAllMarkers() function in Seurat and filtered with (P_adj_ of Wilcoxon’s rank-sum test < 0.05, log_2_ (FC)□>□0.25 and expressed in >25% of cells of the given cluster. E-assembloidE-assembloid

For 10x Genomics data from E-assembloids, *RPS4Y1* expression was used to help determine the source of the cells as the two cell lines used to construct E-assembloids are of different genders. The male AIC-N hESC derivatives were further filtered with *RPS4Y1* expression (normalized and natural-log (log1p) transformed value > 1), the remaining cells were integrated with epiblast or hypoblast derivatives from 3D-cultured embryos. Data integration was performed with IntegrateData() function. Cells from E-assembloids were classified based on 10x Genomics data of 3D-cultured human embryos with the TransferData() function in Seurat, and predicted cell types were displayed on UMAP of integrated Seurat object.

The intercellular communication networks of 3D-cultured human embryos and E-assembloids were analysed following the published method ^76^ implemented in CellChat (https://github.com/sqjin/CellChat), the netVisual_signalingRole function was used to visualize the communication pattern among different cells. VlnPlots of different genes were plotted with VlnPlot function of Seurat R package.

### Statistical analysis

Statistical tests were performed on GraphPad Prism 9 software and Microsoft office Excel 2019. Data were checked for normal distribution and equal variances before each parametric statistical test was performed. Where appropriate, *t*-tests were performed with Welch’s correction if variance between groups was not equal. ANOVA tests were performed with a Dunnett’s multiple comparisons test if variance between groups was not equal. Error bars represent standard deviation in all cases, unless otherwise noted. Figure legends indicate the number of independent experiments and statistical subjects performed in each analysis.

## Notes

### Competing Interest Statement

The authors have declared no competing interest.

